# Modelling the prebiotic origins of regulation & agency in evolving protocell ecologies

**DOI:** 10.1101/2024.11.20.624505

**Authors:** Ben Shirt-Ediss, Arián Ferrero-Fernández, Daniele De Martino, Leonardo Bich, Alvaro Moreno, Kepa Ruiz-Mirazo

**Affiliations:** Donostia International Physics Centre (DIPC – CSIC, UPV/EHU), San Sebastián, Spain; Department of Philosophy, IAS-Research Centre for Life, Mind and Society, University of the Basque Country (UPV/EHU), Donostia-San Sebastián, Spain; Biofisika Institute (CSIC, UPV/EHU), Leioa, Spain; Ikerbasque Foundation, Bilbao, Spain

**Keywords:** Prebiotic systems evolution, minimal metabolism, ecopoiesis, regulation, adaptive agency

## Abstract

How did natural systems develop the first mechanisms of regulation and what for? How could they turn into adaptive agents in a minimal (though deeply meaningful) biological sense? A novel simulation platform, ‘**Araudia**’, has been implemented to address these questions, which are deeply interrelated, in a prebiotic scenario where metabolically diverse protocells are allowed to modify their dynamic behaviour in response to changes in their boundary conditions (e.g., nutrient concentrations in the medium) and/or in the activity of other protocells, including cross-feeding relationships. We extend ‘consumer-resource models’ (CRMs) to a stochastic evolutionary framework in which novelty appears *bottom-up* (i.e., from small changes at the individual protocell level), and a short-term memory may also come forth and spread in the population, to demonstrate that simple adaptive/learning processes can have relevant effects at somatic times (i.e., within the lifetime of single protocells). Our interest in exploring the interplay between metabolic-physiological aspects and ecological-evolutionary ones derives from the fact that it provides a complex causal domain, where various spatial and temporal scales intermingle, and where both the *actual* and the *potential* (pathways/behaviours) must be considered. It is in such a complex domain where the appearance of regulation acquires full significance, as an effective means to navigate phenotypic spaces that become too big for unguided exploration, given the large number of possible functionalities (or functional states) accessible to each proto-cellular agent. The results obtained for several minimalist cases below show that, despite the non-trivial combination of --both endogenous and exogenous--factors involved in the constitution of regulatory mechanisms, these are bound to play a critical role in the survival of infra-biological systems, especially under rapidly changing environmental conditions that longer-term adaptive mechanisms would not be able to cope with.

## Introduction

Life is autonomy in open-ended evolution [Ruiz-Mirazo et al. 2004]. This requires genetically-instructed and cellularly organized metabolisms, whose emergence from physics and chemistry remains a mystery. Our way of tackling this deep scientific riddle favours the early formation of proto-cellular systems, as it was argued for and analysed thoroughly in previous work (see: [Shirt-Ediss et al. 2014; Piedrafita et al. 2017] plus refs. therein). Here we move to a more advanced prebiotic scenario where those relatively simple protocells, which may already display some self-productive and self-reproductive capacities [Ruiz-Mirazo & Mavelli 2008; Mavelli & Ruiz-Mirazo 2013], should transform into increasingly elaborate and efficient metabolic systems. Yet, such systems must have faced a strong bottleneck [Ruiz-Mirazo & Moreno 2024]: the higher their complexity, the larger the space of possible dynamic trajectories/behaviours available to each of them. At earlier stages in the transition from the inert to the living, the main difficulty was probably dealing with the combinatorial explosion in chemical composition and reactivity associated to proto-metabolic networks [Nogal et al. 2023]; nevertheless, once that is overcome and a subset of organic building blocks is selected to construct molecular and organizational complexity, the challenge is to tame what we shall here call “functional expansion” processes. This is not an issue for models of origins of life based on the evolution of self-replicating polymers, since the functional/phenotypic space of those molecular entities stays rather limited (in a first approximation, just two-dimensional: resistance to hydrolysis and replication speed [Wicken 1987; Moreno & Ruiz-Mirazo 2009]), regardless of the amount of possible monomer combinations that one can theoretically calculate, for a given polymer length. However, when one conceives abiogenesis as a process of protocell development [Shirt-Ediss et al. 2017; Ruiz-Mirazo et al. 2020], the individuals of the evolving population consist in compartmentalized micro-reactors, supramolecular entities with a heterogeneous distribution of component parts, whose repertoire of dynamic behaviours (i.e., number of possible stationary states, asymptotic attractors -- or, simply, viable reaction trajectories/configurations) significantly expands as they transform into increasingly complex organizations. Dynamic systems theory strongly supports this idea, having demonstrated, over the years, that the dimensionality and non-linearities present in the description of a system correlate with features like multi-stability, complex behaviour, unexpected bifurcations, etc. (see [Dudkowski et al. 2016] for a nice review on the topic). Furthermore, being open metabolic systems that synthesize their own physical boundaries, protocells must engage, in parallel, in well-controlled exchanges with a variable environment -- if they are to maintain themselves, grow and eventually reproduce under non-equilibrium conditions.

Thus, the way we envision the process of material and organizational complexification towards minimal life involves populations of self-(re-)producing protocells in continuous interaction with their peers and their inert --but variable-- surrounding milieu, as a result of which they develop novel ways of thriving, of operating functionally and adaptively as individuals, even though the effort required is necessarily collective and transgenerational [Ruiz-Mirazo et al. 2024]. This implies modifying both the internal metabolic dynamics of each protocell, as well as the external (competitive and cooperative) relationships that they establish with other members of the population. Therefore, a large *space of possibilities* will be, in principle, accessible to each protocell, even though only a relatively small *subregion of actual trajectories* can be, in practice, probed. Random or unguided searches in such a complex causal context are bound to fail, sooner or later, unless the environment remains relatively stable. But this condition is not realistic in any natural, prebiotic setting -- and less so if each individual is surrounded by similar ones, undergoing chemical transformations and continuous change. Therefore, the chances for these systems to increase their levels of material and organizational complexity strongly depend on the implementation of mechanisms to: (i) search effectively in the space of possible dynamic states/functions available to them, taking into account the inputs and perturbation patterns they get from their environment; and (ii) record and transmit reliably those mechanisms to subsequent generations in the population.

The computational approach and simulation results presented below will focus on the first point (i), assuming that protocells, if they manage to grow, will reproduce without difficulties and transmit to their offspring their composition, properties, the variations they occasionally suffer… as well as what they may “learn” on the way. We are aware that this is a crude simplification, and (ii) --namely, the appearance of hereditary mechanisms-- will also deserve attention in future research, but our primary aim here was to understand how regulation, as a second-order control mechanism [Bich et al. 2016], could contribute to the dynamic robustness of metabolizing protocells when “the possible” becomes much larger than “the actual”. As explained in the next section (Methods and modelling assumptions -- and in the supplementary information, SI), first-order controls would be protocell functional components (oligopeptides -- eventually, proteins) responsible for the uptake and processing of nutrients into byproducts. The modulation of these basic molecular constraints by an additional set of constraints is key for the survival of the protocells in a rapidly varying world where they have too many options at reach. In that context, regulatory mechanisms come to rescue, channelling system responses according to the patterns of change --or the challenges-- that the protocells get from the environment. More specifically, our central interest here will be to investigate the emergence of dynamic behaviours that can be tuned within the lifespan of a single protocell (i.e., ontogenetic or somatic adaptation), even if the process of establishment of the underlying regulatory mechanism, the “assimilation-sedimentation” process itself, may cover much longer, evolutionary timescales (i.e., protocell phylogenies of variable length).

In brief, our approach to the problem consists in offering an evolutionary framework (supported by a new stochastic simulation platform, ‘**Araudia**’) where the complexity of the individuals in the population not only concerns their internal metabolic nature (as compartmentalized, chemical micro-reactors) but is also linked to the proto-ecological relationships that they set up (in particular, cross-feeding), which allow for the co-existence of diverse protocell types within the same environment. For that purpose, our computer platform was built as an elaboration of microbial ‘consumer-resource models’ (CRMs) [Goldford et al. 2018; Marsland et al. 2019; 2020], which try to account for the high levels of species diversity observed in natural ecosystems, moving beyond the classical ‘competitive exclusion principle’ that applies in most evolutionary models (see the critical review [van den Berg et al. 2022] and refs. therein). Keeping metabolically and ecologically distinct species in the pool of evolving protocells is important for our origins-of-life research program, since we also aim to explore the (co-)emergence and evolutionary development of ‘minimal agency’ [Barandiaran et al. 2009; Moreno 2018] within the same prebiotic setting. Indeed, regulatory mechanisms that modulate, specifically, the *outward* behaviour of single protocells (including the interactive dynamics they engage into with their peers) must have constituted a basic requirement for the first adaptive forms of agency to, ever, unfold in nature.

But one step at a time: let us introduce briefly first the main characteristics and assumptions of our computational tool, our stochastic CRM, where occasional changes in the values of the parameters associated to single protocells create “variants” that drive evolution from the bottom-up, so to speak. In addition to several metabolic constraints that make the model thermodynamically consistent, we include a plastic “short-term memory” to channel system responses to environmental challenges by tuning the (rates of change in the) amounts of functional components present in each protocell. This second-order constraint (main novelty of our approach with regard to previous CRMs) will be instrumental to obtain regulatory behaviours of the ‘lac-operon’-kind, as we will report later, in the Results section, together with other findings that involve minimalistic protocell ecologies. Finally, in Discussion and Outlook, we recapitulate our main results, analyse their theoretical implications and suggest future lines of investigation that will expand beyond the rather elementary, “proof-of-principle” cases here explored.

## Methods and main modelling assumptions

Our main effort for this contribution was to encompass, within a single simulation platform, several processes that happen at different spatial and temporal scales: from physical-chemical transformations to protocell metabolisms to ecological networks; and from the on-going (somatic) behaviours of single-protocells to much longer-term, trans-generational trajectories (i.e., evolving phylogenies). This necessarily implied taking some simplifying commitments, which have biassed the research carried out, so it is very important to make them explicit. For starters, entities in our model are of three main types: *molecules* (chemicals: nutrients/byproducts), *protocells* (larger micro-reactors that uptake and process those chemicals through functional components or “enzymes” -- which actually operate as a blend between catalyst and transporter) and *populations* (collections of identical protocells, each corresponding to a ‘sub-species’ biomass).^1^ As depicted in Fig. 1, the amounts of all of these entities change in time, stochastically, within a well-stirred chemostat (a flow-reactor with a fixed global volume).

**Figure 1.**
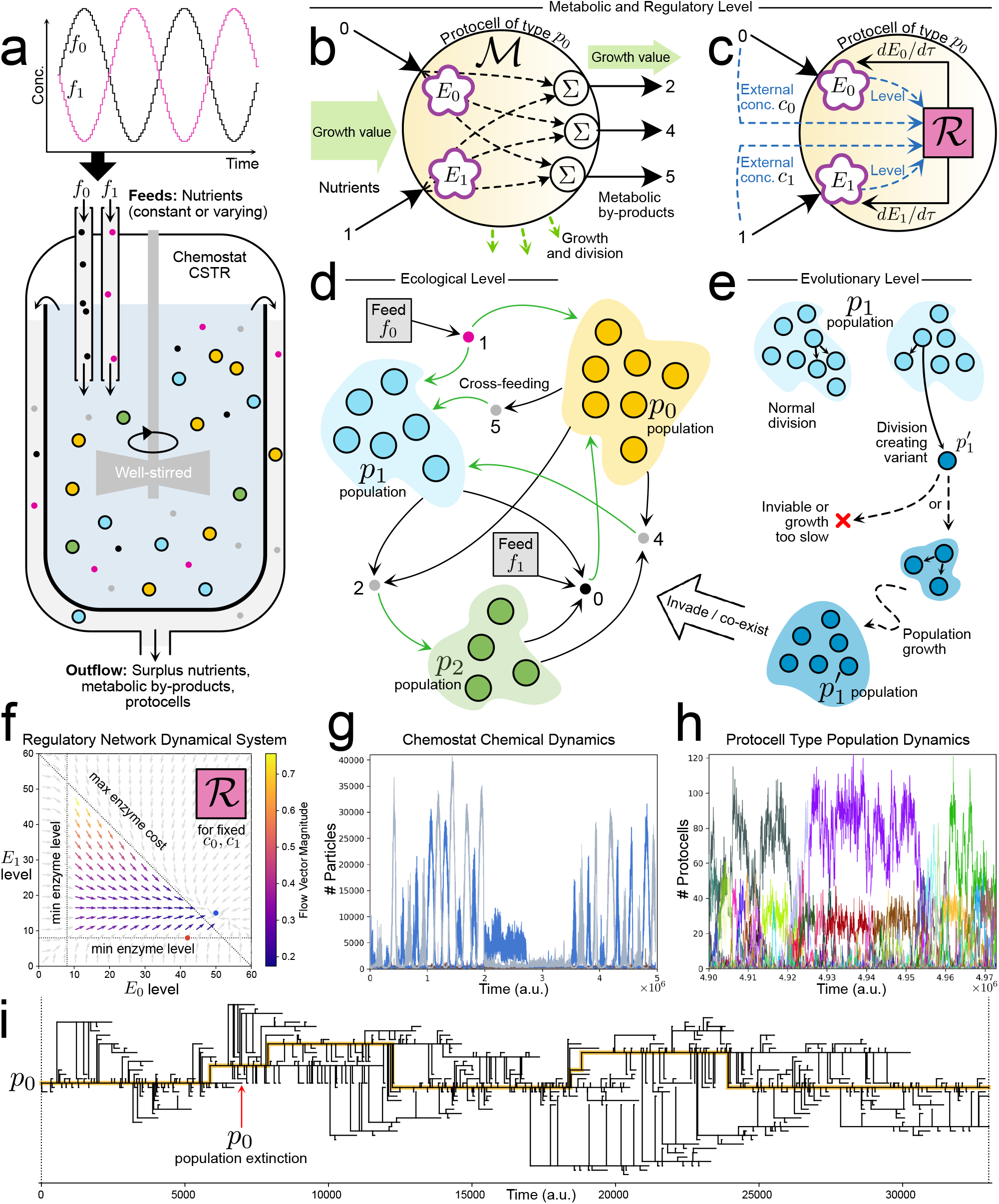
Modified consumer-resource model to study the emergence of minimal regulation strategies in an evolving ecology of protocells. **(a)** Protocell population dynamics take place in a well-stirred constant volume chemostat with varying nutrient inputs. Each protocell has **(b)** a metabolic network and **(c)** an (initially disabled) regulatory network (see text). **(d)** Populations of different protocell types can co-exist in the flow reactor by using strategies such as cross-feeding. **(e)** Further, the current protocell ecology can be disrupted by evolutionary events whereby a variant protocell type is created and grows as a new population. Remaining panels show example dynamics from simulations: **(f)** Enzyme-level dynamics of a regulatory network (see text); **(g)** Nutrient and metabolic by-product chemical dynamics in the reactor; **(h)** Zoom of protocell population dynamics in the reactor; **(i)** Lineage tree showing the evolution of one protocell type in the initial condition, for a short time.

On top of the Gillespie [1976; 1977] algorithm implementing the basic reaction kinetics, our computational platform (‘**Araudia**’ -- described in more detail in the SI) introduces the possibility of evolution by modifying the value of some parameter and creating, in this way, a protocell “variant”. Such evolutionary changes are only allowed, with a certain probability, at division steps (which, in turn, only occur if there is sustained protocell growth) and will scale up to the population level (or not), becoming a significant piece of the biomass (or not), depending on how well the new protocell variant does in the chemical context it encounters at birth, and in the interactive dynamics it establishes with other (co-existing) protocell sub-species. If a protocell variant eventually gives rise to another variant that remains within the “protocell phylogeny trunk” (yellow line in Fig. 1.i) the former will be considered a *significant* variant, since its contribution to the protocell lineage is long-term. Hence a standard simulation begins with a number of protocells (typically, in the hundreds) of one or several species and, provided that they thrive, a “cloud of sub-species” will be subsequently generated. All this involves quite rich and complex dynamics, reminiscent of the ‘quasi-species’ model [Eigen & Schuster 1979], even if the underlying processes and assumptions here are completely different. Our individuals grow and divide into identical daughters, without errors, unless a variant appears, but they do not behave like “sequence replicators”: rather, they constitute compartmentalized chemical bioreactors (more precisely, the linear sum of the metabolic pathways that define each protocell -- see Fig. 1b) evolving under variable external conditions and under the “selective pressure” of being directly washed out, with a certain probability, from the tank (i.e., no explicit fitness function will be employed, just a global ‘reactor flow speed’ or ‘dilution rate’ – more details in the SI).

The rate at which a protocell of a given type/sub-species thrives is determined by the amount of “growth value” it is effectively importing from external nutrients, minus a cost for “self-maintenance”, as well as a “global resource cost” that applies to the total number of functional components that uptake and process those nutrients (i.e., the first-order controls or “enzymes” present in each protocell -- one kind for each nutrient in the diet, so a ‘trade-off’ is forced, following [Posfai et al. 2017]). In order to ensure physical/ thermodynamic consistency, we also add the rule that the overall rate of (nutrient) growth value flowing into any metabolism operating in the chemostat should be strictly higher than the overall rate of (byproduct) growth value flowing out of it. Thus, in the deterministic limit, our model corresponds to a standard miCRM (microbial consumer-resource model) that implements an ‘OR’ function for nutrients (i.e., protocells can feed on one or all nutrient inputs in their diet, and nutrient uptake rates are essentially independent of each other). However, all of this turns quite more complicated by the introduction of an additional, novel feature: a “memory function” that operates as a flexible (second-order) constraint on the dynamics of each protocell type (i.e., affecting the nutrient preferences of that particular protocell type). Since this is a key aspect that makes our work distinct from previous CRMs, let us explain it a bit further.

With the aim to investigate the origin of regulatory mechanisms and their impact on the behaviour and adaptive capacities of proto-cellular systems, we searched for a plastic function that could correlate protocell states with changes/perturbations coming from the external medium -- beyond their intrinsic potential to respond to those changes through random variations, across evolutionary time scales. In other words, we wanted to include a possible source of “ontogenetic plasticity” for the protocells, in addition to their inherent “phylogenetic adaptivity” (as self-reproducing, evolving systems). On these lines, instead of adding the possibility of modifying directly the number of functional components of a protocell, we thought that it was more interesting to modulate the *rate of change* of those components, so we used, for that purpose, a “neural ODE”, following [Chen et al. 2018]. The neural ODE (a network with variable weights, whose outputs are the values of the corresponding derivatives -- see Fig. 1c and Fig. 1f), can be active or inactive, depending on the situation under analysis, but it is never explicitly trained: it is just “tinkered” through the actual evolutionary trajectories obtained in our simulations. So what we have implemented here has nothing to do, in practice, with a standard neural network or a machine learning algorithm: it is just an abstract formulation of the high-order constraint (i.e., a correlation function) that can serve as a “memory” for our protocells, if they manage to take advantage of it (which is not always the case).^2^

Let us describe, next, what this theoretical approximation to the problem has provided in terms of simulation results. All the data and plots for analysis reported below come from computer simulations carried out with ‘**Araudia**’, which were run in an HPC cluster [DIPC Hyperion]. Parameter settings can be found in the SI material.

## Results

The stochastic dynamics of three different minimal protocell ecologies were here investigated, searching specifically for protocells that spontaneously evolve functional “lac-operon”-like regulatory networks as a way of enhancing their survival: i.e., regulatory networks that can quickly and flexibly switch the nutrient preference of a host protocell to match the most abundant nutrient in the environment. In this prebiotic context, the capacity for ‘nutrient preference switching’ can be seen as a rudimentary agent property (see Discussion section, below), and one which can be explored, anyway, in the non-equilibrium and well-stirred conditions assumed in our model.

### General conditions for our ‘in silico chemostat’ experiments

Two nutrient chemicals were supplied to the flow reactor, either both at a constant level (at 200 concentration units each), or each varying as a sine wave, 180 degrees out of phase with the other (sine waves between 40 and 200 concentration units). We tested two main cases: a −REG control case, where protocells could not develop regulatory networks, and a +REG case, where protocells started with disabled regulatory networks but had the potential to turn functional through “evolutionary tinkering” of the neural ODE weights. In the −REG control case, the only way that protocells could adapt to changing nutrient abundance was by using phylogenetic adaptation (the birth of fitter, more suited variants) to modify the levels of their import enzymes and metabolic network weights, over relatively long time scales. All results reported below measure enzyme dynamics across the entire evolutionary lineage of protocells beginning with the protocell type(s) in the initial condition, as illustrated by the yellow line in Figure 1(i).

Our analysis typically focuses on internal enzyme dynamics, because they directly reflect the nutrient preferences of protocells. Although the evolving protocell system also has other interesting aspects, such as protocell population dynamics, reactor chemical dynamics, or the evolution of metabolic network weights, these are secondary to our aims here and are not explicitly reported. Finally, in all simulations reported here, major evolutionary innovations were disabled and, therefore, each protocell variant imported the same nutrients and excreted the same metabolic by-products as its parent type. Nevertheless, loosely speaking, “ecologies” of different protocell types or protocell ‘sub-species’, each with its corresponding biomass (see Footnote 1), are typically generated throughout our simulations. The full mathematical model description and detailed initial conditions are given in the SI material.

### Case Study 1: ‘single protocell species’

The first and simplest scenario we investigated was the evolution of a population of a single protocell species ‘p0’ that could feed off either the supplied nutrients 0 or 1, or both of them (Figure 2a). Each of the supplied nutrients had equal growth value. Protocell type p0 already possessed a passive ‘null strategy’ to survive in changing nutrient abundance conditions: it could import and grow from the nutrient present at the time, and simply pay the maintenance cost for the import enzyme for the non-present nutrient. However, evolutionary development of a regulatory network could enable a more sophisticated and efficient response: namely, dynamic allocation of the limited import enzyme budget toward the currently most abundant nutrient in the environment.

**Figure 2:**
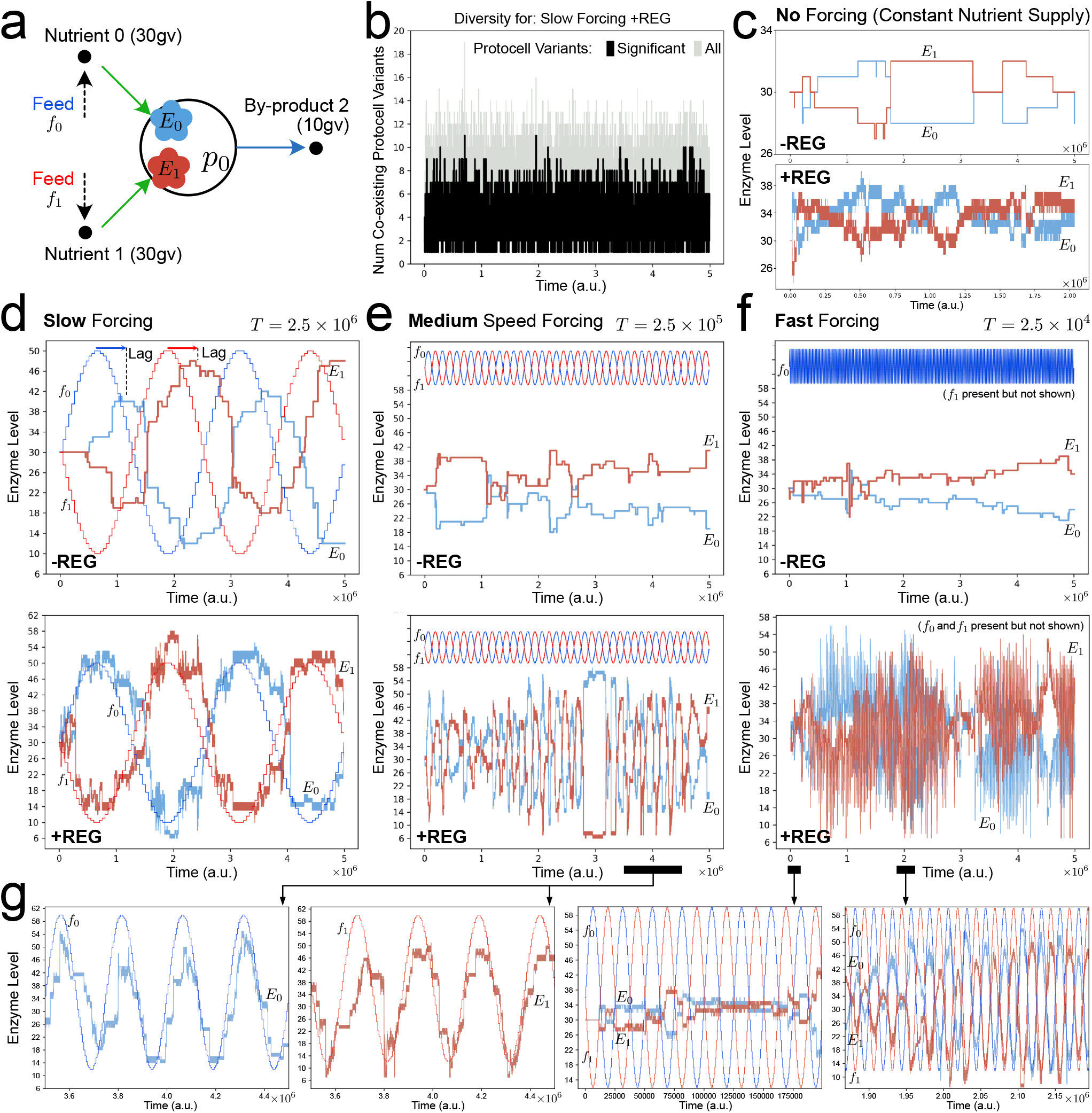
Dynamics of internal enzyme levels across an evolutionary lineage stemming from a single protocell type, when protocells are deficient in regulatory networks (−REG) and when protocells have the potential to develop functional regulatory networks (+REG), under constant, slow, medium and fast-varying nutrient forcing. **(a)** Reactor starts with a population (n=100) of protocell variant p0 which eats nutrient 0 or nutrient 1, or both to grow. **(b)** Many variants of p0 arise and co-exist during the course of evolution. ‘Significant’ protocell variants are those which give rise to further variants. **(c)** Under constant nutrient input (reactor feeds f0 = f1 = 200 conc. units) enzyme levels across the evolutionary lineage remain equally split or diverge symmetrically in both −REG and +REG cases. Panels **(d-f)** show enzyme dynamics across the evolutionary lineage when reactor feeds f0 and f1 now vary as sine waves, between 40 and 200 conc. units, 180 degrees out of phase. When sinusoidal forcing is slow, −REG protocells can still slowly adapt via phylogenetic changes (d, upper figure), but faster nutrient forcing prohibits phylogenetic adaptation (e, f, upper figures). Conversely, under faster nutrient forcing, +REG protocells evolve “lac-operon”-like dynamics, allocating (limited) internal enzyme levels in phase with the varying abundance of nutrients in the environment (e,f, lower figures). **(g)** Zoom plots of regulatory network enzyme dynamics in response to nutrient forcing. Note that nutrient forcing sine waves f0 and f1 use a separate y-axis that is not shown.

When the reactor feeds supplied constant nutrient levels (at 200 concentration units each), both in the −REG and +REG cases, the E0 and E1 enzyme levels both either evolve to hover around the midpoint of 30 units (Figure 2c), or they diverge symmetrically from the midpoint. This divergence can be rationalised by the fact that nutrients 0 and 1 had equal growth values, and under constant nutrient supply, they could be imported almost interchangeably. As we just advanced above, even in this simple case of a single protocell species, in which the population is under constant nutrient supply, the initial biomass (∼100 identical protocells) systematically evolves to diverse pieces of biomass over time: that is, from a single protocell type in the initial condition to a mean of around 4 protocell types or ‘quasi-species’ at any one instant (Figure 2b).

If nutrient abundance is varied as a slow sine wave with period T = 2.5×10^6^ time units, enzyme levels across the evolutionary lineage of protocell types now track the forcing, both in the −REG and +REG cases (Figure 2d). In the −REG case, more of a lag occurred between the change in abundance of nutrient in the reactor feed and the subsequent change in internal import enzyme level. This can be rationalised by noting that p0 has a shorter phenotype size (of 4) in the −REG control condition, which leads to a decrease frequency of variants and hence a longer ‘wait time’ for adaptation. Conversely, p0 in the +REG case has a greater internal complexity, reflected by a longer phenotype size (45), and this yields an increased frequency of variants. Additionally, after such a “functional expansion”, in the longer phenotype there exist many regulatory network parameters which can subtly affect enzyme levels. These latter two factors allow less lagged adaptation to the varying nutrient source in the +REG case. It is important to remark that in the+REG case (under constant or slowly changing nutrient concentrations), the adaptation is still purely phylogenetic: the regulatory network is not functionally modifying the dynamics of the enzyme levels. This is evidenced in Figure 4a(i) where a +REG protocell type at the end of the evolutionary lineage is tested against nutrient forcing, but with evolutionary variants disabled. In this case, the regulatory network does not dynamically respond to the nutrient forcing.

Under medium speed forcing of nutrient abundance (sine wave period T = 2.5×10^5^ time units), effective phylogenetic adaptation is no longer possible in the −REG case (Figure 2e, top). For the simulation parameters used, in the −REG condition, ∼25000 time steps are required to produce a significant variant (SITable 4), giving only 10 significant variants per sine wave period. This is insufficient to properly adapt enzyme levels to the oscillation of nutrient abundance, hence enzyme levels evolve the null strategy as in the case of no forcing (Figure 2c). Conversely, in the +REG case under medium speed forcing, the behaviour is markedly different and there are frequent bursts of successful adaptation to the varying nutrients (Figure 2e, bottom). For the simulation parameters used, in the +REG condition, ∼5000 time steps are required to produce a significant variant (SI Table 4), yielding on average 50 variants per sine wave period. Adaptation to the nutrient source forcing in this case is a combination of both phylogenetic adaptation and partial emergence of functional regulatory networks. Figure 4a(ii) shows that a late-stage protocell type has evolved partially functional regulatory network enzyme dynamics when evolutionary variants are disabled. In this example, the regulatory network adapts internal enzymes E0 and E1 to follow external nutrient concentrations when [c0]>[c1], but not the other way around.

Finally, under fast nutrient forcing (sine wave period T = 2.5×10^4^ time units), the −REG condition is again unable to utilise phylogenetic adaptation and enzyme levels follow the null strategy, as in Figure 2e (top). However, in the +REG case (Figure 2f, bottom), fast nutrient forcing causes the protocell type to evolve a functional regulatory network. As there are only 5 significant variants born per nutrient oscillation under fast forcing (on average), phylogenetic adaptation is limited. As evidenced in Figure 4a(iii), evolution tinkers a regulatory network able to autonomously change import enzyme levels, over short time scales, in phase with the reactor nutrient forcing. It is interesting to note that even when an efficient regulatory solution has evolved, sometimes, due to the complexity of the selection process, this solution can be temporarily outcompeted by a protocell type with a non-functional regulatory network. Thus, the evolution of functional regulatory networks has a punctuated or bursting behaviour (consistent with a rugged fitness landscape). This is seen both in the medium (Figure 2e, bottom) and fast speed (Figure 2f, bottom) forcing cases.

### Case Study 2: ‘minimal mutualism’

We also investigated the evolutionary emergence of regulation mechanisms in a minimal mutualistic protocell ecology (Figure 3a). The mutual ecology already has an intrinsic robustness or ‘null strategy’ to deal with changing nutrient concentrations: the symmetric cross-feeding relationships meant that when nutrient 0 was abundant p1 could cross-feed from p0 via metabolic by-product 2, and when nutrient 1 was abundant, p0 could cross-feed from p1 via metabolic by-product 3, instead. Our aim was to see if regulatory networks would evolve to implement a more efficient, dynamic response whereby E1 in p0 and E2 in p1 would increase in synchrony as nutrient 0 became abundant, while E1 in p1 and E3 in p0 would increase in synchrony as nutrient 1 became abundant. Under constant nutrient inputs (Figure 3b), for both −REG and +REG cases, the cross-feeding ecology evolves the strategy wherein p0 mainly feeds from nutrient 0 (by up−REGulating E0), and p1 mainly feeds from nutrient 1 (by up−REGulating E1). The REG+ case arrives faster at this strategy because of the increased frequency with which variants are produced. During evolution, again several quasi-species develop around the original two protocell types.

**Figure 3.**
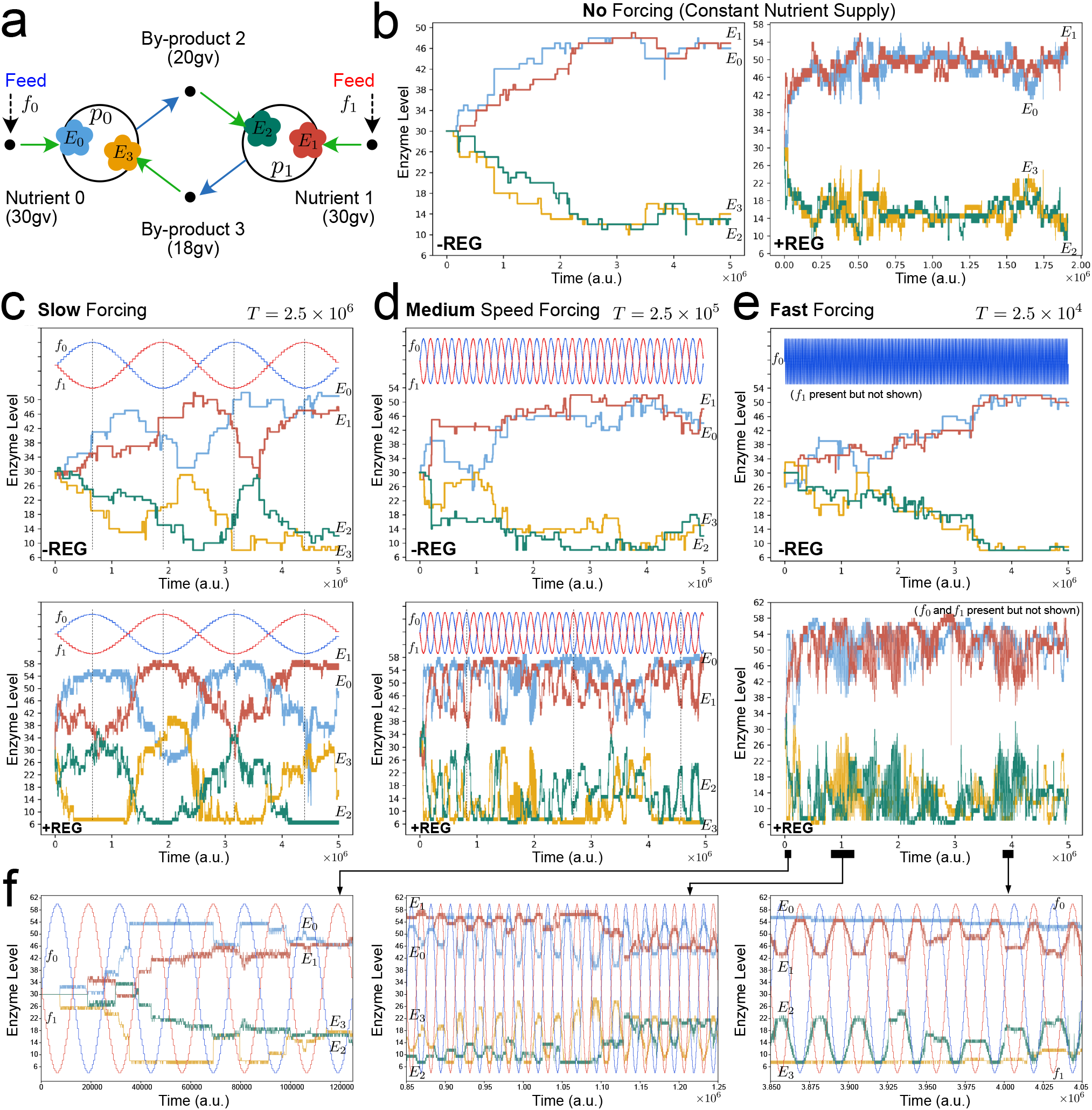
Dynamics of internal enzyme levels across the evolutionary lineage of a mutualistic ecology consisting of two protocell types, without (−REG) and with (+REG) potential to develop regulation mechanisms, under different nutrient forcing conditions. **(a)** Reactor starts with two protocell populations (n=100 each) that cross-feed from each other. Enzyme dynamics across the evolutionary lineage under: **(b)** constant nutrient supply (f0 = f1 = 200 conc. units); **(c)** slow sinusoidal nutrient forcing where nutrients 0 and 1 oscillate between 40 and 200 conc. units, and are and supplied 180 degrees out of phase to the reactor; **(d)** medium speed sinusoidal nutrient forcing; **(e)** fast sinusoidal nutrient forcing. **(f)** Zoom plots of enzyme dynamics in panel (e). Note that on all plots, nutrient forcing sine waves f0 and f1 use a separate y-axis that is not shown.

**Figure 4.**
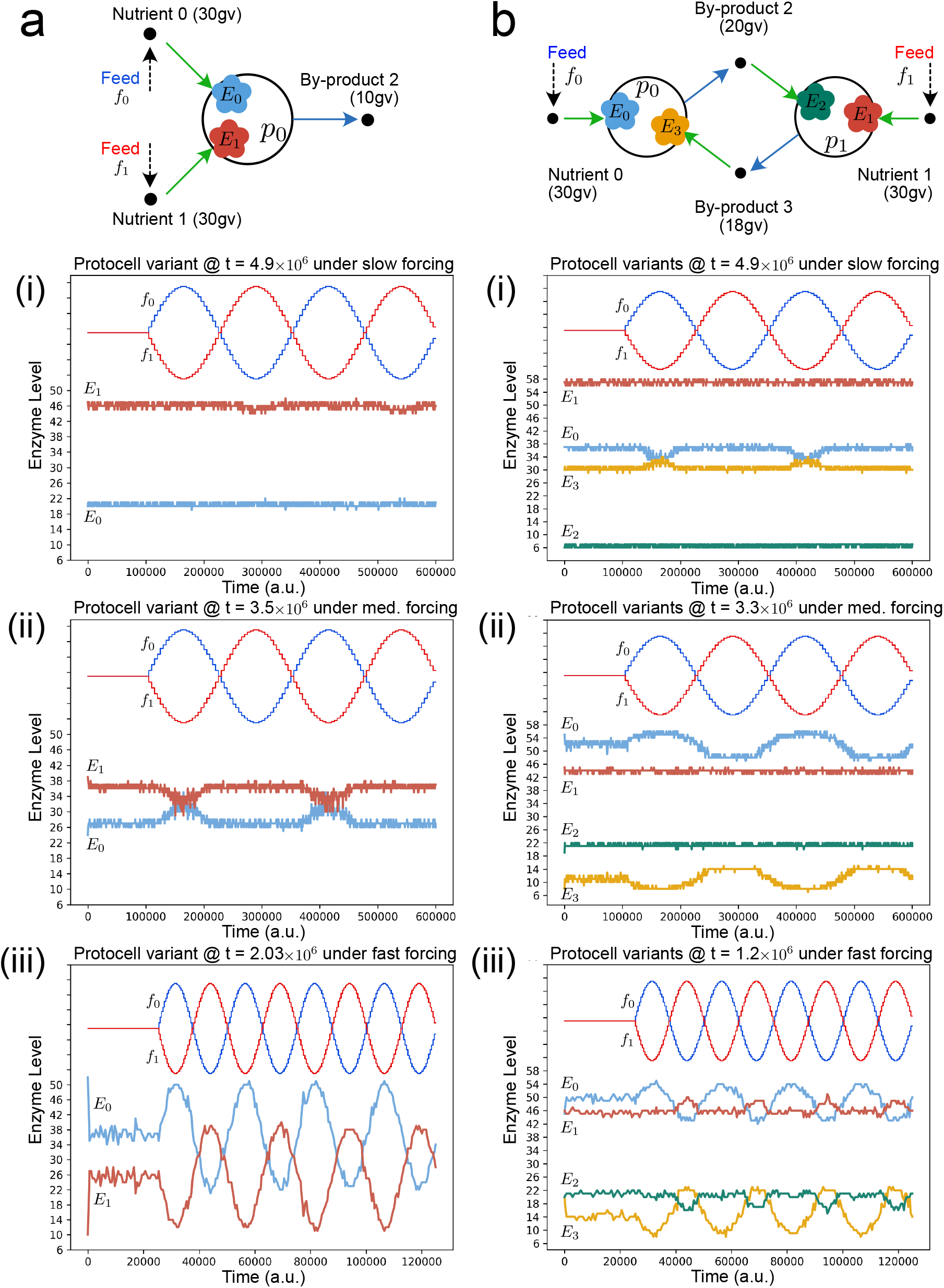
Dynamic responses of evolved regulatory networks in protocell types isolated from the main evolutionary lineage in simulations from Figures 2 and 3. Protocell types are added to a new reactor in 100 copy number and exposed to nutrient cycling. Evolutionary mutations over time are disabled (no phylogenetic adaptation) and thus the source of enzyme level changes originates solely from regulatory networks, exposing what they have ‘learned’. **(a)** For the single protocell type simulations in Figure 2. **(b)** For the two protocell type ecology simulations in Figure 3.

When nutrients vary as two slow sine waves in anti-phase (Figure 3c), internal enzyme levels in the protocell types across the evolutionary lineage oscillate in-phase with nutrient availability (+REG), or in-phase but lagged (−REG), similar to the Case Study 1. Figure 4b(i) shows that phylogenetic adaptation is mainly responsible for the enzyme level changes, with a regulatory response that is, in fact, not adequate (it brings down E0 levels when nutrient 0 is present in the medium). Increasing nutrient abundance to a medium oscillation speed (Figure 3d) demonstrates that phylogenetic adaptation is insufficient to change enzyme levels in the −REG case, and enzyme levels over the evolutionary lineage become similar to the no-forcing case (Figure 3b). In the +REG case, enzyme levels do track nutrient abundance, and this is caused by a combination of phylogenetic adaptation and short-term dynamics of the regulatory networks (Figure 4b(ii)).

Using a fast oscillation speed for nutrient abundance in the +REG case (Figure 3e, bottom) causes bursts of enzyme level activity in-phase with nutrient abundance. This activity is mainly controlled by regulatory networks (Figure 4b(iii)) as phylogenetic adaptation is weak. Interesting to note is that under medium and fast forcing, p0 and p1 don’t always change their enzyme levels in synchrony: there are times when p0 is active alone, and other times when p1 is active alone, and times when both are inactive. The ‘extreme regulation’ strategy (whereby E0 and E2 are at their maximum levels, while E1 and E3 are at their minimum, when nutrient 0 is abundant, and vice-versa when nutrient 1 is abundant) does not evolve because nutrient uptake rates also depend on local chemical concentrations in the reactor, which diminish as uptake rates get higher.

### Case Study 3: ‘binary asymmetric ecology’

As a final case, we investigated a protocell ecology that featured both cross-feeding and competition (Figure 5a). With potential for competition, survival of this ecology was much more sensitive to initial conditions as compared to Case Studies 1 or 2. We found that starting with particular metabolic weights (like those listed in Figure 5a), this ecology could sustain when evolving under no nutrient forcing (Figure 5b), and under slow nutrient forcing (Figure 5c,d,e), but not under medium speed or fast nutrient oscillations. As such, regulatory networks with functional dynamics did not evolve in this ecological case. However, the larger phenotypic space in the +REG case still allowed this ecology to exhibit another interesting effect. Starting from identical initial conditions, depending on stochastic fluctuations, this ecology could show two qualitatively different modes of phylogenetic adaptation. Either the p0 lineage adapted to the slow nutrient oscillations and the p1 lineage always cross-fed (Figure 5d), or the p0 lineage did not adapt and instead the p1 lineage selectively cross-fed, adapting to the oscillating abundance of metabolic by-product 2. These different modes of adaptation were accompanied by distinctive population dynamics.

**Figure 5.**
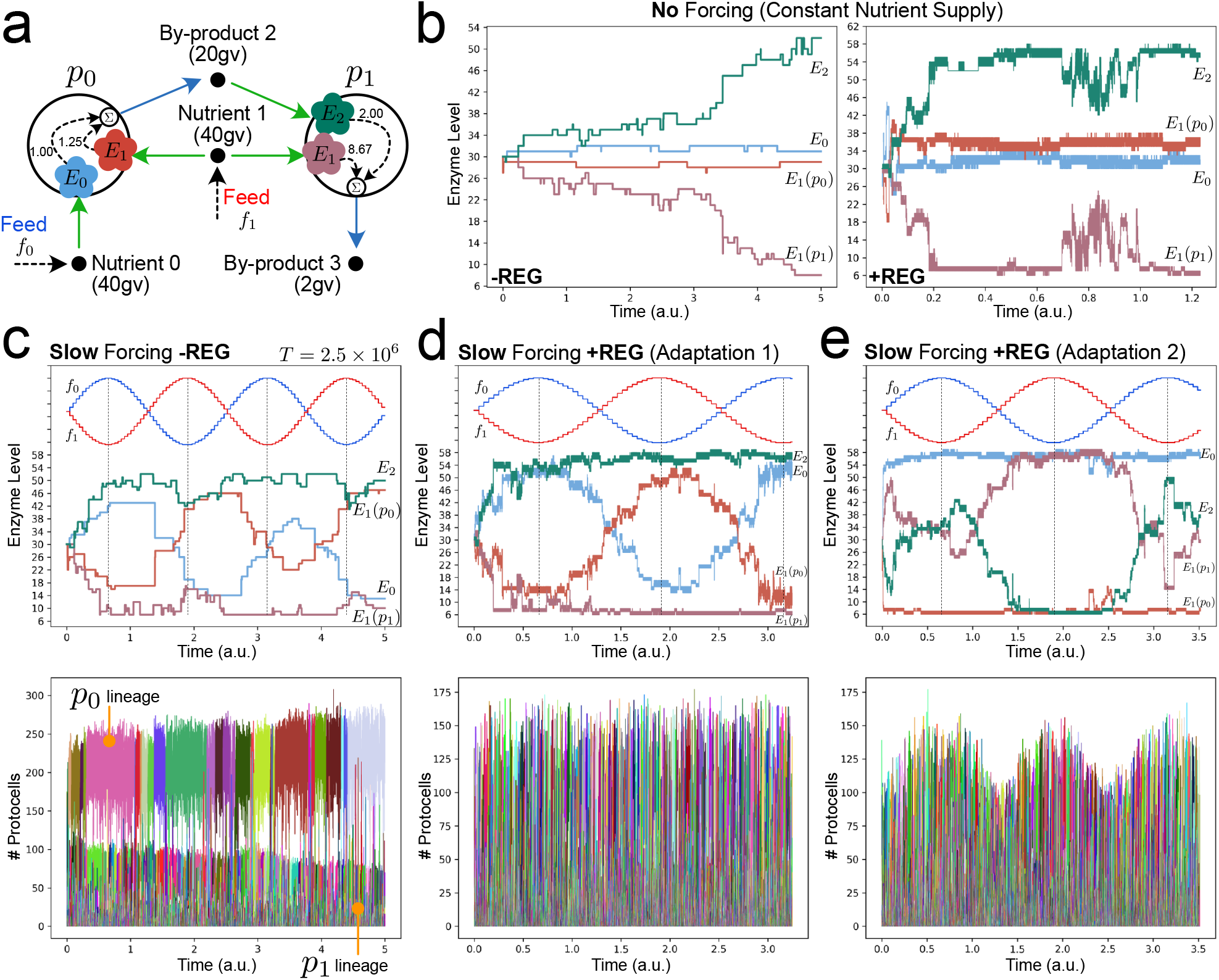
Different modes of phylogenetic adaptation of a co-operative/competitive minimal protocell ecology. **(a)** Reactor starts with two protocell populations (n=100 each) that can cross-feed and/or compete for supplied nutrient 1. Long-term survival of this ecology under nutrient forcing and evolutionary mutations is highly sensitive to the initial conditions used (unlike ecologies in Figures 2 and 3). Initial metabolic weights used for all simulations are shown. Enzyme dynamics across the evolutionary lineage under: **(b)** constant nutrient supply (f0 = f1 = 200 conc. units); **(c, d, e)** slow sinusoidal nutrient forcing. When protocells have the potential to develop regulatory mechanisms (+REG) under slow forcing conditions, then qualitatively different modes of phylogenetic adaptation can be achieved from the same initial condition (as shown in **(d)** and **(e)**). Both modes have distinct population dynamics. For the parameters used, functional regulatory networks did not develop in this ecology. Further, the ecology could not survive medium speed or fast forcing conditions.

## Discussion & Outlook

There are several aspects worth highlighting from the result just reported, as a way of recapitulating our main findings and addressing then the limitations of what we have explored so far, together with future avenues of research that could be tackled with ‘**Araudia**’. First of all, a note on the complexity of the dynamics involved in our simulations. Nothing should be taken for granted, really, as the last case study has shown us. One could have expected, from the results gathered in the first (‘single protocell species’) and second (‘minimal mutualism’) scenarios, that regulatory behaviour was going to emerge, also in the third (‘binary asymmetric ecology’), as the most adequate strategy to face environmental perturbations, especially when their frequency is increased. But this was not obvious. Of course, more simulation runs and initial conditions/parameters should be tried (including some comparative analysis with other theoretical approaches that could be addressing similar problems: e.g., [Tikhonov & Monasson 2017; Advani et al. 2018; Altieri & Franz 2019; Pham & Kaneko 2024]). Yet, the diverse, non-linear combination of factors and scales within our computational approach makes predictability a real issue. Similarly, the degree of success of somatic/ontogenetic adaptive mechanisms (in comparison to evolutionary/phylogenetic ones -- and also in conjunction with the relative frequency of external oscillations) is not easy to guess, despite the apparent general trend observed in the first two cases: i.e., the more stringent the environmental forcing, the more relevant regulatory behaviour seemed to turn. Still, in the third case, under rapidly varying conditions, regulation did not show up in time (provoking the actual collapse of one of the protocell species).

Another point to be remarked is the fact that evolution and regulation are not disconnected from each other but, rather, asymmetrically coupled dimensions of the same phenomenon: namely, the transformation of the stochastic dynamic behaviour of protocell populations at longer-term and wider scales. Of course, this somehow derives from our theoretical premises and modelling assumptions, but it was solidly confirmed by the simulation results obtained so far. Evolution (together with the presence of an external variation pattern “susceptible to assimilation” by the protocells -- irregular modifications of the boundary conditions would be much harder to deal with) is necessary for the emergence of regulation; in turn, the appearance of regulatory mechanisms modifies the evolutionary potential of the system (and not just in somatic terms but also in a strictly phylogenetic sense -- as it was plain clear in the last case study, where the option for regulation, although it was a possibility, did not fructify). That is due to the increase in phenotypic space and, thereby, in the possibilities for mutation available to protocells. Thus, when regulatory mechanisms start operating in a system/population, the corresponding functional space is expanded and, with that, also evolvability per se. Indeed, the question is not just how big the number of state variables and parameters^3^ gets, but the amount of possible states that becomes accessible when all of them are put together. Effective regulation (i.e., the active use of the flexible correlation function added to the algorithm) plays a double role in this context: it expands *the space of the possible* and, in the same move, it provides new means to explore that space, enabling protocells to choose *actual* pathways/behaviours that happen to be better solutions to the challenges they face.

Nevertheless, the search for eventual stationary states, dynamic attractors or just optimized solutions where all the relevant factors (metabolic, ecological and evolutionary) are consistently integrated is not straight-forward, given the number of components, constraints and non-trivial, multi-scale causal loops we attempt to capture within the same modelling framework. When this project was started, we considered making things even harder for the algorithm, but those initial ideas will have to wait for future implementations. For instance, the plastic “memory function” has been tried, so far, without any additional production/maintenance cost, which is not realistic: any material subsystem devoted to assimilation/ detection and regulatory tasks should take a significant part of the energy budget of a protocell (even if this is relatively smaller than the production/maintenance of other, more basic, functional components of the system). Strong simplifications were also made regarding the metabolic functioning of each protocell, whose real intricacies may have important evolutionary and ecological implications (see, e.g.: [Tomaiuolo et al. 2008], [Mori et al. 2016], or [Meijer et al. 2020]). Likewise, spatial constraints do not play much of a role in our well-stirred, flow-reactor setting, although we are perfectly aware of their relevance, not only at the level of the individual (which is something that we have worked on, previously, from different approximations [Piedrafita et al. 2017; Lauber et al. 2023]), but also in the collective and interactive domain (for instance, in the emergence of cross-feeding relationships [Meijer et al. 2020]).

An obvious extension of the work reported here, which has been mainly focused on micro-evolutionary dynamics and minimalistic ecological cases, would involve a higher degree of ‘protocell species diversity’ (i.e., more heterogeneity in terms of metabolic ‘ways of life’, or ‘diets’, in the population) and, thus, also the development of more complicated protocell-protocell relationships. ‘**Araudia**’ already presents the simulation options and features required for that research, so the potential is clearly there, but it has not been exploited yet. Even though we have tried scenarios with various diets/species, these were kept constant in most of our simulations runs to date. Yet, the plan is to explore more complex ecological settings, either involving a bigger number of protocell species already in our initial conditions, or allowing for their appearance during the simulation runs -- to monitor, then, whether the corresponding “niches” (new biomass pieces) remain interconnected and dynamically/structurally stable, thereafter, or not. Our strategy here has been to begin with simple (binary) cases, but it could be case that the competitive exclusion principle operates more strongly in those scenarios (which would be –partly-- the reason for the results obtained in our third case study): indeed, there is evidence that multispecies microbial communities emerge and remain stable when a minimal threshold of diversity is crossed [Chang et al. 2023].

On these lines, various kinds of changes in the way protocell diets are implemented will be tried in upcoming work: (i) make possible, in the course of our simulations, the modification in the nutrients and/or byproducts of any given protocell metabolism in the population; (ii) explore “co-consumption” scenarios, in which secreted compounds are re-utilized as nutrients by the same species [Millard et al. 2023] -- a possibility that was ruled out for our current simulation runs; (iii) transform the ‘OR’ nutrition function (which we used by default, up to now) into a more demanding ‘AND’ (which would apply to other specific situations -- e.g., carbon vs. nitrogen bearing substrate consumption [Kleijn et al. 2023]); (iv) let other intermediary compounds play additional roles in protocell-protocell interactions (e.g., appearance of toxins); (v) include the option of secreting/detecting compounds to/in the environment that are not being used as nutrients/metabolites with a direct fitness cost (e.g., indirect ‘signals’ or chemical cues [Pacheco et al. 2019]). All this will surely open a whole new world of possibilities, where protocells unfold novel behaviours in terms of competition/cooperation strategies with their peers. Even if protocell capacities to switch/modulate their nutrient preferences, as they were studied here, may already be taken as rudimentary forms of agency, it is in this wider context where the emergence of prebiotic *adaptive* agents could be more fruitfully addressed, because “speciation events” (or “macroevolutionary transitions”, in our terminology) should transform protocell interactive features more deeply -- making them, at the same time, subject to regulation.

In conclusion, we can say that the theoretical and computational modelling work presented in this article has provided interesting insights into the problem that we were initially addressing, especially by demonstrating that protocells can “learn” from their interactions with the environment and perform actual regulatory behaviours that operate in somatic times, making them, in effect, more robust. Through that effort, we may have contributed to unveil some aspects of the rather intricate link between regulation and evolution in the natural world, whereas the goal of connecting regulation with the prebiotic evolution of more sophisticated forms of agency, on the way towards proper biological agency, awaits future endeavours.

## Supporting information

Supplementary Information (Mathematical Model)

## Acknowledgments

This work has been supported by the John Templeton Foundation (Science of Purpose -- Grant No. 62220), as well as by grant PID2023-147251NB-I00 for project ‘Outagencies’, and grant PID2023-146408NB-I00, for project ‘Cellular Phase Transitions’, funded by MCIN/AEI/10.13039/501100011033 and by ERDF/EU. In addition, the IAS-Research group members (see affiliations above) also acknowledge funding from the Basque Government (Grupos Consolidados: Ref. IT1668-22), while AF-F, DDM and KR-M are grateful to the Fundación Biofisika Bizkaia (FBB -- Basque Excellence Research Centre -- UPV/EHU & CSIC) for continuous support. We thank the technical staff at DIPC for support in using the Hyperion HPC cluster.

## Authors’ Contributions

BS-E, LB, AM and KR-M conceived the research line. BS-E developed and implemented the CRM model using an optimised code, ran most simulations and analysed their results. DDM contributed to the technical specification of the platform, and AF-F to obtaining and analysing additional simulation data. BS-E produced all Figures and elaborated the SI, as well as a first draft of the Results section. KR-M wrote a first draft of the rest of the manuscript. All authors contributed with constructive feedback to the final (submitted) version of both main text and SI.

## Supplementary Information

Supplementary Information containing the mathematical model description accompanies this article.

## Competing Interests

All authors declare that they have no competing interests.

## Footnotes

1 The term ‘sub-species’ is the adequate one here, because we keep the ‘species’ category for a (larger) group of protocells that share a common “diet” or “ecological niche” (i.e., the same set of nutrients/byproducts in the chemostat, regardless of the values of all other simulation parameters). Although our computational platform allows for “speciation events” (i.e., the stochastic appearance of protocell variants that imply a change of diet), we are not reporting any simulation results that include such events yet. In other words, we are focusing, for the time being, on “micro-evolution” dynamics. Nevertheless, we would like to keep this terminology, in view of future expansions of the current work towards the exploration of “macro-evolutionary” transitions.

2 This also helps us circumvent the problem of proposing a more specific “recording mechanism”, which would have to be highly speculative, given the current state of affairs in the field of prebiotic evolution.

3 Our modelling approach can be interpreted as a ‘constructive dynamical system’, where state variables (protocell population levels, reactor chemical levels, enzyme levels) change over time under fixed parameters but, at longer temporal scales, evolution can also change parameters (metabolic network weights, regulatory network weights etc.), as well as the actual number of state variables (e.g., when new protocell populations arise). In this way, the evolutionary algorithm transforms the structure of the (stochastic) dynamical system itself, throughout the simulation.

## Notes

### Competing Interest Statement

The authors have declared no competing interest.

