## Supplementary Information (Mathematical Model) for "Modelling the prebiotic origins of regulation & agency in evolving protocell ecologies"

### **Araudia**: A stochastic simulation platform to study the origins of regulation and agency in evolving (proto)cell ecologies

Ben Shirt-Ediss[1], Arián Ferrero-Fernández[2,3], Daniele De Martino[3,4], Leonardo Bich[2], Alvaro Moreno[1] and Kepa Ruiz-Mirazo[2,3⋆]

[1]Donostia International Physics Centre (DIPC – CSIC, UPV/EHU), San Sebastián, Spain

[2]Department of Philosophy, IAS-Research Centre for Life, Mind and Society, University of the Basque Country (UPV/EHU), San Sebastián, Spain

[3]Biofisika Institute (CSIC, UPV/EHU), Leioa, Spain

[4]Ikerbasque Foundation, Bilbao, Spain

#### Contents

**Supplementary Note 1**
**Model Notation** **2**

**Supplementary Note 2**
**Stochastic Model Formulation** **3**

**Supplementary Note 3**
**Deterministic Model Formulation** **8**

**Supplementary Note 4**
**Regulatory Network** **9**

**Supplementary Note 5**
**Evolution Operator** **16**

**Supplementary Note 6**
**Parameter Values for Simulations in Paper** **18**

**Supplementary Note 7**
**Lineage Sizes and Variant Numbers for Simulations in Paper** **20**

**Supplementary Note 8**
**Computational Optimisations** **21**

### Supplementary Note 1 Model Notation

#### Units

- Units are coloured in **teal**
- Time is in abstract unit $\tau$, volume in abstract unit $V$, concentration in abstract unit $C = V^{-1}$

#### Rates

- $\Gamma$ denotes event rates (unit $\tau^{-1}$)
- $\gamma$ denotes concentration rates (unit $C\tau^{-1}$)

#### Concentrations, Particle Numbers, Enzyme Levels

- $c_i$ and $C_i$ are the concentration and particle number of chemical species $i$ respectively
- $p_\sigma$ and $P_\sigma$ are the concentration and particle number of protocell type $\sigma$ population, respectively. A protocell 'type' corresponds to a given protocell 'subspecies': see Footnote 1 of the main paper text.
- $E_{\sigma i}$ is the level of the functional component (or 'enzyme', as a shortcut term) importing/processing nutrient $i$ in all individuals of protocell type $\sigma$ population

#### Sets

- $N_\sigma$ is the set of chemical species $i$ that protocell type $\sigma$ consumes as nutrients
- $B_\sigma$ is the set of chemical species $i$ that protocell type $\sigma$ leaks as metabolic by-products
- $N_i$ is the set of protocell types $\sigma$ that import chemical species $i$ as a nutrient
- $B_i$ is the set of protocell types $\sigma$ that leak chemical species $i$ as a metabolic by-product

If a protocell type imports a chemical as a nutrient, it cannot excrete the same chemical as a metabolic by-product, and vice-versa:

$$N_\sigma \cap B_\sigma = \emptyset \qquad N_i \cap B_i = \emptyset \tag{1}$$

This excludes the possibility of a by-product becoming a nutrient for the same protocell type at a later stage. We note this is a strong assumption: e.g. *E.coli* can take in sugars and secrete acids, only to take in those acids at a later stage [S1].

### Supplementary Note 2 Stochastic Model Formulation

Supplementary Figure 1 summarises the 10 fundamental transition types in the stochastic model. Individual transitions always involve incrementing/decrementing a single particle number, protocell population number, or enzyme level in the system. The rate of each transition (in events per time unit) is defined below. The model is executed using the Gillespie SSA algorithm [S2].

#### Flow Reactor (Chemostat) Transition Rates

The rate that particles of chemical species $i$ enter the (well-stirred) reactor vessel when present at concentration $f_i$ (unit $C$) in the reactor feed is given by:

$$\Gamma_{\mathrm{i}}^{\mathrm{inflow}} = Q_{\mathrm{in}} f_i \qquad = \mu \Omega f_i \quad \text{unit } \tau^{-1} (\text{particles per unit time}) \tag{2}$$

where parameter $\mu = \frac{Q_{\mathrm{in}}}{\Omega}$ (unit $\tau^{-1}$) is the dilution rate, i.e. the fraction of the flow reactor volume $\Omega$ (unit $V$) displaced by incoming solvent per unit time. $Q_{\mathrm{in}}$ (unit $V\tau^{-1}$) denotes the number of solvent volume units entering the reactor per unit time. Quantity $\frac{1}{\mu}$ (unit $\tau$) is the 'mean residence time' of the reactor. Reactor feed concentrations $f_i$ can be constant over time or varied as a step function (e.g. stepped sine wave, or square wave) defined in a feed schedule CSV file. Using a step function rather than a continuous function allows simulation of the system as a consecutive series of autonomous dynamical systems, rather than as a non-autonomous dynamical system.

The rate that chemical species $i$ washes out of the reactor vessel when present inside at $C_i$ copies is:

$$\Gamma_{\mathrm{i}}^{\mathrm{washout}} = \mu C_i \quad \text{unit } \tau^{-1} (\text{particles per unit time}) \tag{3}$$

The rate at which protocell type $\sigma$ washes out of the reactor vessel when present inside at $P_\sigma$ copies is:

$$\Gamma_{\sigma}^{\mathrm{washout}} = \mu P_\sigma \quad \text{unit } \tau^{-1} (\text{protocells per unit time}) \tag{4}$$

Note that no protocells flow into the reactor via the reactor feeds.

#### Single Protocell Transition Rates

The rate at which a single protocell of type $\sigma$ imports nutrient chemical $i$ depends on the level of internal import enzyme $E_{\sigma i}$ and on the concentration of the nutrient $c_i = C_i/\Omega$ in the reactor vessel via Monod kinetics:

$$\Gamma_{\sigma i}^{\mathrm{nutrient}} = E_{\sigma i} \frac{c_i}{1 + c_i} \quad \text{unit } \tau^{-1} (\text{particles per unit time}) \tag{5}$$

where we assume $K_m = 1$ for all enzymes (i.e. all enzymes are equally effective at importing and processing nutrients).

The enzyme level $E_{\sigma i}$ (unit $\tau^{-1}$) is measured as the number of particles the enzyme imports into the protocell per unit time when external nutrient concentration $c_i$ is saturating.

The rate that a single protocell of type $\sigma$ leaks a metabolic by-product chemical of species $i$ back into the reactor vessel is simply a weighted sum of the current import rates of all nutrients:

$$\Gamma_{\sigma i}^{\mathrm{byproduct}} = \sum_{n \in N_\sigma} \Gamma_{\sigma n}^{\mathrm{nutrient}} M_{ni}^{\sigma} \quad \text{unit } \tau^{-1} (\text{particles per unit time}) \tag{6}$$

| | Rate | Eq. | Description | State Change |
|---|---|---|---|---|
| T1 | $\Gamma_{i}^{inflow}$ | 2 | Reactor inflow of 1 copy of chemical species $i$ | +1 copy of chemical species $i$ |
| T2 | $\Gamma_{i}^{washout}$ | 3 | Reactor wash out of 1 copy of chemical species $i$ | -1 copy of chemical species $i$ |
| T3 | $\Gamma_{\sigma}^{washout}$ | 4 | Reactor wash out of 1 copy of protocell type $\sigma$ | -1 copy of protocell type $\sigma$ |
| T4 | $\Gamma_{\sigma i}^{nutrient}$ | 5 | Protocell of type $\sigma$ import 1 copy of nutrient species $i$ | -1 copy of chemical species $i$ |
| T5 | $\Gamma_{\sigma i}^{byproduct}$ | 6 | Protocell of type $\sigma$ leak 1 copy of by-product species $i$ | +1 copy of chemical species $i$ |
| T6 | $\Gamma_{\sigma}^{divide}$ | 14 | Protocell of type $\sigma$ divide with 2 identical daughters | +1 copy of protocell type $\sigma$ |
| T7 | $\Gamma_{\sigma}^{unviable}$ | 16 | Protocell of type $\sigma$ becomes unviable | -1 copy of protocell type $\sigma$ |
| T8 | $\Gamma_{\sigma}^{mutant}$ | 15 | Protocell of type $\sigma$ divide with 1 daughter mutated | **New variant protocell type added** |
| T9 | $\Gamma_{\sigma i}^{enzyme+}$ | 26 | Protocell of type $\sigma$ internal enzyme $i$ increase 1 unit | +1 unit of enzyme $i$ in protocell type $\sigma$ |
| T10 | $\Gamma_{\sigma i}^{enzyme-}$ | 27 | Protocell of type $\sigma$ internal enzyme $i$ decrease 1 unit | -1 unit of enzyme $i$ in protocell type $\sigma$ |

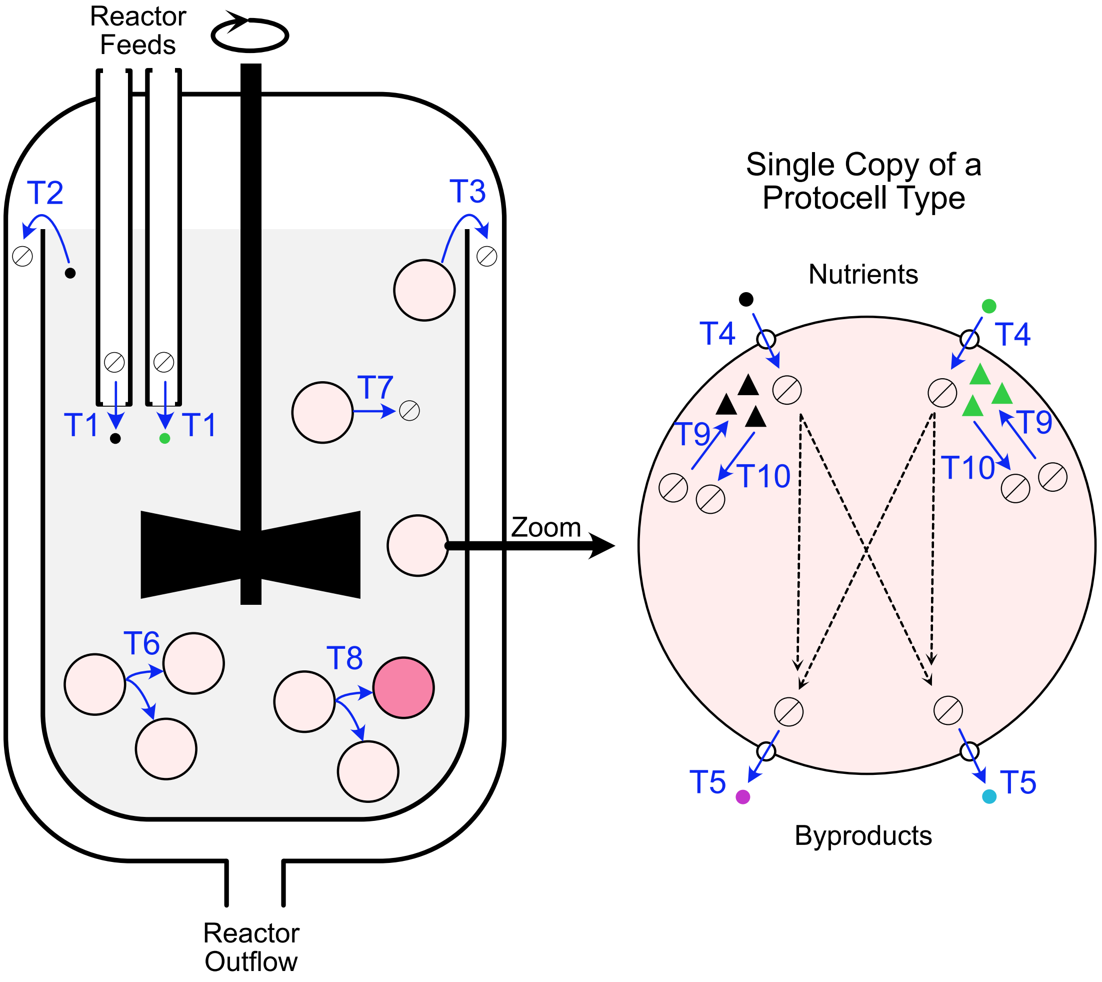


Supplementary Figure 1: Stochastic transition types and diagram showing where transition types happen in chemostat/protocells.

where $M^{\sigma}_{ni}$ is the metabolic matrix (following Ref. [S3]). Matrix elements are zero or positive and can be viewed as metabolic 'weights'. Weight $(n, i)$ where $n \neq i$ denotes how many molecules of by-product $i$ are leaked following the import of a single nutrient molecule $n$. Weight values $< 1$ denote that more than 1 nutrient molecule must be imported before a single by-product molecule is leaked. Note that the summation term in Eq. 6 implies that a protocell can grow on *any or all* of the available nutrients (an OR dependency). However, nutrient preference can be modified by the regulatory network (see Supplementary Note 4).

By determining the production rates of by-products, the metabolic matrix also determines how much "growth value" can be extracted from each nutrient particle imported into a protocell:

$$\Delta v_{\sigma n} = v_n - \sum_{b \in B_\sigma} v_b M^{\sigma}_{nb} \quad \text{unit } v \tag{7}$$

where $\Delta v_{\sigma n}$ and $v_n$ are the "useable" and total growth values of a single nutrient particle of species $n \in N_\sigma$, respectively. Value $v_n$ is defined in the Chemical Universe, see Supplementary Note 6.

To ensure a basic thermodynamic consistency that prevents spontaneous energy/mass creation in protocell ecologies, following Ref. [S3] the weights in the metabolic matrix must obey:

$$v_n > \sum_{b \in B_\sigma} v_b M^{\sigma}_{nb} \quad \text{unit } v \tag{8}$$

for each nutrient $n \in N_\sigma$. That is, $\Delta v_{\sigma n} > 0$. This ensures that, over a defined time, the total growth value flowing into a protocell (as nutrients) is always greater than the total growth value leaked out of a protocell (as metabolic by-products).

We further assert that a protocell type excretes all metabolic by-products at non-zero rates. This can be seen as a simplification of the fact that in metabolism, there always exists a division between catabolic and anabolic pathways, the former providing for energy and always secreting something. This requirement also promotes cross-feeding between different protocell types. We constrain metabolic matrix weights such that a minimum fraction $\beta$ of the growth value of each imported nutrient is always leaked back to the environment as metabolic by-products / wastes:

$$\frac{v_n - \Delta v_{\sigma n}}{v_n} \geq \beta \quad \text{dimensionless} \tag{9}$$

The total growth value per unit time available for protocell growth (leading to eventual division) is given by:

$$\lambda_\sigma = \left( \sum_{n \in N_\sigma} \Gamma^{\text{nutrient}}_{\sigma n} \Delta v_{\sigma n} \right) - R^{\text{maintain}} - R^{\text{enzymes}}_{\sigma} \quad \text{unit } v\tau^{-1} \tag{10}$$

That is, the rate of import of each nutrient multiplied by the useable growth value per nutrient particle, minus a fixed resource cost for protocell maintenance $R^{\text{maintain}}$ (unit $v\tau^{-1}$) and a resource cost for maintaining the nutrient import/processing enzymes at their current levels:

$$R^{\text{enzymes}}_{\sigma} = \sum_{n \in N_\sigma} E_{\sigma n} e_n \quad \text{unit } v\tau^{-1} \tag{11}$$

where $E_{\sigma n}$ is the level of the internal enzyme importing nutrient $n$ and $e_n$ (unit $v$) is the growth value cost for maintaining one unit of the latter enzyme. Value $e_n$ is defined in the Chemical Universe, see Supplementary Note 6.

Protocells are systems of finite size and finite metabolic capacity and thus internal enzyme levels cannot be infinite. Following Ref. [S4] we introduce a maximum resource cost for enzymes, forcing a trade-off in enzyme levels:

$$R_\sigma^{\text{enzymes}} \leq R_{\text{MAX}}^{\text{enzymes}} \quad \text{unit } \upsilon\tau^{-1} \tag{12}$$

This constraint is the same for all protocell types. Parameter $R_{\text{MAX}}^{\text{enzymes}}$ determines the upper limit rate that a protocell type can import nutrients from the reactor environment. Additionally, we introduce a minimum level that each import enzyme must have in a protocell type:

$$E_{\sigma i} \geq E_{\text{MIN}} > 0 \quad \text{unit } \tau^{-1} \tag{13}$$

This assumes there is some baseline 'constitutive' expression of all enzymes, and was found necessary to prevent some active regulatory networks from killing their host species.

The rate at which a single protocell divides into two identical daughters is proportional to the positive magnitude of $\lambda_\sigma$:

$$\Gamma_\sigma^{\text{divide}} = \max(0, g\lambda_\sigma) \quad \text{unit } \tau^{-1}(\text{new protocells per unit time}) \tag{14}$$

where parameter $g > 0$ (unit $\upsilon^{-1}$) converts growth value per unit time into protocell divisions per unit time. Every protocell type has the same value $g$ parameter.

The rate at which a single protocell divides with *one* of the two daughters harbouring random mutations is 1 every $\mathcal{E}_{\text{div}}$ divisions, on average (again independent of protocell type):

$$\Gamma_\sigma^{\text{mutant}} = \frac{1}{\mathcal{E}_{\text{div}}}\Gamma_\sigma^{\text{divide}} \quad \text{unit } \tau^{-1}(\text{mutant protocells per unit time}) \tag{15}$$

The rate that a single protocell becomes metabolically unviable (a form of disappearance different to being washed out of the reactor) is proportional to the negative magnitude of $\lambda_\sigma$:

$$\Gamma_\sigma^{\text{unviable}} = \max(0, -g\lambda_\sigma) \quad \text{unit } \tau^{-1}(\text{unviable protocells per unit time}) \tag{16}$$

Finally, the rates of change for internal enzyme levels inside a protocell ($\Gamma_{\sigma i}^{\text{enzyme+}}$ and $\Gamma_{\sigma i}^{\text{enzyme-}}$ for each enzyme importing nutrient species $i$) are determined by the regulatory network and are given in Supplementary Note 4.

#### Protocell Population Transition Rates

Transition rates used in the simulation are at the level of protocell type populations. Because all protocells in a population of protocell type $\sigma$ are considered identical, the conversion from individual protocell rates (given above) to population rate simply entails multiplying by population size $P_\sigma$. Note that this conversion does not apply to enzyme rates $\Gamma_{\sigma i}^{\text{enzyme+}}$ and $\Gamma_{\sigma i}^{\text{enzyme-}}$ because these are set by the regulatory network of a protocell type (see Supplementary Note 4) and all members of the protocell type have the same internal enzyme state.

#### State Vector has Dynamic Size

The model is a *constructive* dynamical system in the sense that, in addition to state changing over time, the number of state variables and their coupling can additionally evolve over time:

- System state vector size decreases by 1 when a protocell type goes extinct, e.g. $P_\sigma = 0$. Note that population extinction is not, in itself, an event. When all populations go extinct then the simulation is stopped.
- System state vector size increases by 1 when a $\Gamma_\sigma^{\text{mutant}}$ event fires. The new variant daughter (modified from the parent protocell type using the evolution operator, see Supplementary Note 5) is added to the state vector with a copy number of 1.

#### Initial Condition

The initial condition of the model is specified by:

- Parameters (Supplementary Note 6)
- Chemical Universe (Supplementary Note 6)
- Reactor feed forcing schedule
- The initial ecology of protocell types:
  - Which protocell types are present (each type has metabolic weights, import enzyme levels and regulatory network weights as free parameters)
  - The copy number of each protocell type

At time $t = 0$ the reactor starts at full solvent volume $\Omega$ with all chemical concentrations in the reactor vessel equilibrated to the reactor feed concentrations. The initial protocell ecology is added to this state. Each protocell type is at 100 copies. After a "burn in" time (typically 1000-10000 time steps) ecology population levels and reactor vessel chemical concentrations become properly equilibrated. From this point, the simulation begins.

### Supplementary Note 3 Deterministic Model Formulation

The stochastic model is required to conduct the full simulation. It properly captures: (i) events sensitive to chance fluctuations, like population amplifications and (ii) protocell mutations/extinctions which require tracking single copy numbers. The Gillespie algorithm can also naturally handle the changing state space dimension of the system caused by new variants/extinctions.

However, a deterministic model can be applied in parallel, to *verify* the implementation of the stochastic model. It can be run on successive simulation 'segments' where in each segment (i) the protocell ecology has a fixed number of protocell types (system state space size is fixed) and (ii) the reactor feeds (step functions) are not changing in concentration. The deterministic model agrees exactly with the stochastic model in the thermodynamic limit (very large system size), but it is often a useful check also at practical system sizes. The deterministic model (omitting the regulatory network part) is similar in formulation to the Microbial Consumer-Resource Model [S3, S5].

Because reactor volume $\Omega$ is always constant, concentrations and particle numbers are related by:

$$c_i = \frac{C_i}{\Omega} \qquad p_\sigma = \frac{P_\sigma}{\Omega} \quad \text{unit } C \tag{17}$$

Reactor volume $\Omega$ effectively specifies the number of particles per abstract concentration unit.

#### ODE Set for Concentration of Each Chemical in Reactor Vessel (Resources)

$$\frac{dc_i}{d\tau} = \mu\left(f_i - c_i\right) - \gamma_i^{\text{nutrient}} + \gamma_i^{\text{byproduct}} \quad \text{unit } C\tau^{-1} \tag{18}$$

where:

$$\gamma_i^{\text{nutrient}} = \frac{1}{\Omega}\sum_{\sigma\in N_i} P_\sigma \Gamma_{\sigma i}^{\text{nutrient}} \qquad = \sum_{\sigma\in N_i} p_\sigma \Gamma_{\sigma i}^{\text{nutrient}} \quad \text{unit } C\tau^{-1} \tag{19}$$

and:

$$\gamma_i^{\text{byproduct}} = \frac{1}{\Omega}\sum_{\sigma\in B_i} P_\sigma \Gamma_{\sigma i}^{\text{byproduct}} \qquad = \sum_{\sigma\in B_i} p_\sigma \Gamma_{\sigma i}^{\text{byproduct}} \quad \text{unit } C\tau^{-1} \tag{20}$$

#### ODE Set for Concentration of Each Protocell Type in Reactor Vessel (Consumers)

$$\frac{dp_\sigma}{d\tau} = \gamma_\sigma^{\text{grow}} - \mu p_\sigma \quad \text{unit } C\tau^{-1} \tag{21}$$

where:

$$\gamma_\sigma^{\text{grow}} = \frac{1}{\Omega} P_\sigma g \lambda_\sigma \qquad = p_\sigma g \lambda_\sigma \quad \text{unit } C\tau^{-1} \tag{22}$$

#### ODE Set for Enzyme Level in Each Protocell Type

Enzyme level derivatives are calculated by Eq. 23 in Supplementary Note 4. In the deterministic model, different to the stochastic model, internal enzyme levels are treated as *continuous* quantities.

#### Supplementary Note 4 Regulatory Network

Regulatory networks inside protocells are implemented as a modified type of Neural ODE set [S6]. The instantaneous rate of change of the *mean* level of enzyme $E_{\sigma i}$ (in protocell type $\sigma$) importing nutrient $i$ is described by:

$$\frac{1}{k_\tau}\frac{dE_{\sigma i}}{d\tau} = \mathrm{NN}^i_\sigma\left([\mathbf{E}_\sigma, \mathbf{c}_\sigma]\right) - k_{\sigma i}E_{\sigma i} - \xi^{\mathrm{MAX}}_{\sigma i} + \xi^{\mathrm{MIN}}_{\sigma i} \quad \text{unit } \tau^{-1}\text{(levels per unit time)} \tag{23}$$

where:

$$\xi^{\mathrm{MAX}}_{\sigma i} = E_{\sigma i} \times b_1 \times \max\left(0, \frac{R^{\mathrm{enzymes}}_\sigma}{R^{\mathrm{enzymes}}_{\mathrm{MAX}}} - 1\right) \quad \text{unit } \tau^{-1}\text{(levels per unit time)} \tag{24}$$

$$\xi^{\mathrm{MIN}}_{\sigma i} = b_2 \times \max\left(0, \frac{E_{\mathrm{MIN}}}{E_{\sigma i}} - 1\right) \quad \text{unit } \tau^{-1}\text{(levels per unit time)} \tag{25}$$

In summary:

- The protocell regulatory network is a dynamical system where (i) internal enzyme levels are state variables $\mathbf{E}_\sigma = [E_{\sigma 1}, E_{\sigma 2}, \ldots]$ and (ii) the external concentrations of nutrients imported by the enzymes $\mathbf{c}_\sigma = [c_1, c_2, \ldots]$ are boundary conditions (themselves variable, but instead influenced by population dynamics in the reactor).

- The neural network term $\mathrm{NN}^i_\sigma([\mathbf{E}_\sigma, \mathbf{c}_\sigma])$ determines the *production* rate of the enzyme $E_{\sigma i}$, given a neural network input vector consisting of row vector $\mathbf{E}_\sigma$ concatenated to row vector $\mathbf{c}_\sigma$. Potentially, $\mathrm{NN}^i_\sigma([\mathbf{E}_\sigma, \mathbf{c}_\sigma])$ can be a very complex function of the input vector, determined by a weight set specific to protocell type $\sigma$.

- The first order decay term $k_{\sigma i}E_{\sigma i}$ determines the decay rate of import enzyme $i$ based on its abundance.

- The $\xi$ terms are conditional functions to ensure that the regulatory network always produces enzyme level dynamics which obey the constraints in Eq. 12 and Eq. 13. They activate only when enzyme levels go out of bounds:
  - When the current total enzyme cost is too high i.e. $R^{\mathrm{enzymes}}_\sigma > R^{\mathrm{enzymes}}_{\mathrm{MAX}}$, then term $\xi^{\mathrm{MAX}}_{\sigma i}$ makes *all* enzyme derivatives more negative to push back enzyme levels into the viable region. The magnitude of 'push back' each enzyme level experiences depends on the enzyme level $E_{\sigma i}$ and multiplier $b_1$. Zero level enzymes cannot turn negative (although $E_{\mathrm{MIN}}$ makes it unlikely that enzyme levels get to zero).
  - When an individual enzyme drops below the minimum level allowed i.e. $E_{\sigma i} < E_{\mathrm{MIN}}$, then term $\xi^{\mathrm{MIN}}_{\sigma i}$ boosts the derivative for that particular enzyme such that it crosses back into the viable region. The magnitude of boost depends on multiplier $b_2$. Constants $E_{\mathrm{MIN}} > 0$ and $b_2 > 0$ are set sufficiently high, such that enzyme levels cannot reach 0 (avoiding a division by zero condition in Eq. 25).
  - $\xi^{\mathrm{MAX}}$ can be interpreted as the inability of the regulatory network to allocate resources that cannot be paid for metabolically. $\xi^{\mathrm{MIN}}$ can be interpreted as a baseline level expression of enzymes.

- Parameter $k_\tau > 0$ controls the overall response time scale of the regulatory network (see below). This parameter is the same for all protocell types.

- Weight $k_{\sigma i}$ (first-order enzyme decay constant) controls the steady state level of the enzyme $E^*_{\sigma i}$ when the neural network has output 0.5 (see below).

Important notes:

- Enzyme degradation terms $k_{\sigma i}E_{\sigma i}$ and $\xi^{\mathrm{MAX}}_{\sigma i}$ are both multiplied by the current enzyme level, which means that once at zero, enzyme levels cannot turn negative (but, they can increase in the positive direction).
- Splitting the enzyme dynamics into a neural network production term (bounded between 0 and 1) and a decay term ensures that the regulatory network dynamical system has *at least one steady state* for each enzyme (see below).
- All protocells in a population of protocell type $\sigma$ have the same regulatory network in the same enzyme level state.

#### Stochastic Rates of Enzyme Level Changes

The rate at which the level of enzyme importing nutrient species $i$ increases by 1 unit inside a protocell of type $\sigma$ is given by:

$$\Gamma^{\text{enzyme+}}_{\sigma i} = \max(0, \frac{dE_{\sigma i}}{d\tau}) \quad \tau^{-1}(\text{enzyme levels per unit time}) \tag{26}$$

The rate that the same enzyme level decreases by 1 unit is:

$$\Gamma^{\text{enzyme-}}_{\sigma i} = \max(0, -\frac{dE_{\sigma i}}{d\tau}) \quad \tau^{-1}(\text{enzyme levels per unit time}) \tag{27}$$

We note that the true rates are $\Gamma^{\text{enzyme+}}_{\sigma i} = k_\tau \left(\mathrm{NN}^i_\sigma\left([\mathbf{E}_\sigma, \mathbf{c}_\sigma]\right) + \xi^{\mathrm{MIN}}_{\sigma i}\right)$ and $\Gamma^{\text{enzyme-}}_{\sigma i} = k_\tau \left(k_{\sigma i}E_{\sigma i} + \xi^{\mathrm{MAX}}_{\sigma i}\right)$. However, for computational tractability, we set the rates to reflect **only** the nett changes in enzyme levels, to reduce the number of events firing.

#### Example Regulatory Network Importing Single Nutrient

Supplementary Figure 2 illustrates how the import enzyme derivative is calculated in the simplest regulatory network in a protocell type $\sigma$ importing a single nutrient. Also, it defines the neural network weight matrices referenced below. Supplementary Figure 3 shows the types of dynamic behaviour this simplest regulatory network can produce.

#### Calculation of Neural Network Term

Calculation of the neural network term in Eq. 23 involves propagating inputs through a multi-layered perceptron in three steps described below.

1. Entries in the input vector $[\mathbf{E}_\sigma, \mathbf{c}_\sigma] = [E_{\sigma 1}, E_{\sigma 2}, \ldots, c_1, c_2, \ldots]$ are first normalised into a $[0, 1]$ interval by performing:

$$\overline{E_{\sigma i}} = \frac{E_{\sigma i}}{E_{\mathrm{MAX}}} \qquad \overline{c_i} = \frac{c_i}{f_i^{MAX}} \tag{28}$$

where:

$$E_{\mathrm{MAX}} = \frac{R^{\text{enzymes}}_{\mathrm{MAX}}}{\min(e_i)} \tag{29}$$

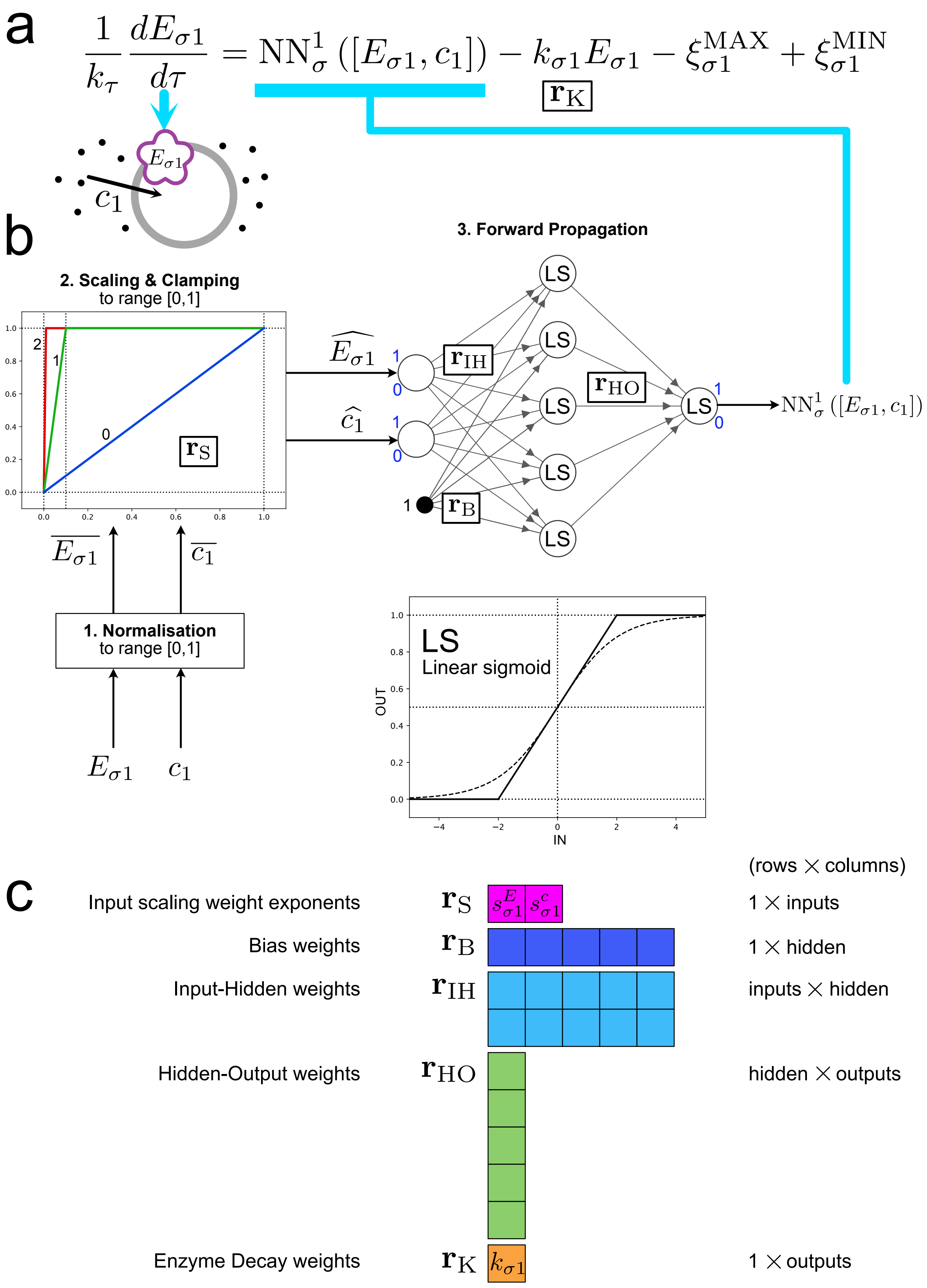


Supplementary Figure 2: Regulatory network for a protocell type $\sigma$ importing a single nutrient $c_1$. (a) Equation describing dynamics of import enzyme $E_{\sigma 1}$. (b) Illustration of neural network term determining $E_{\sigma 1}$ production rate. (c) Weight matrices of the regulatory network (23 weights in total).

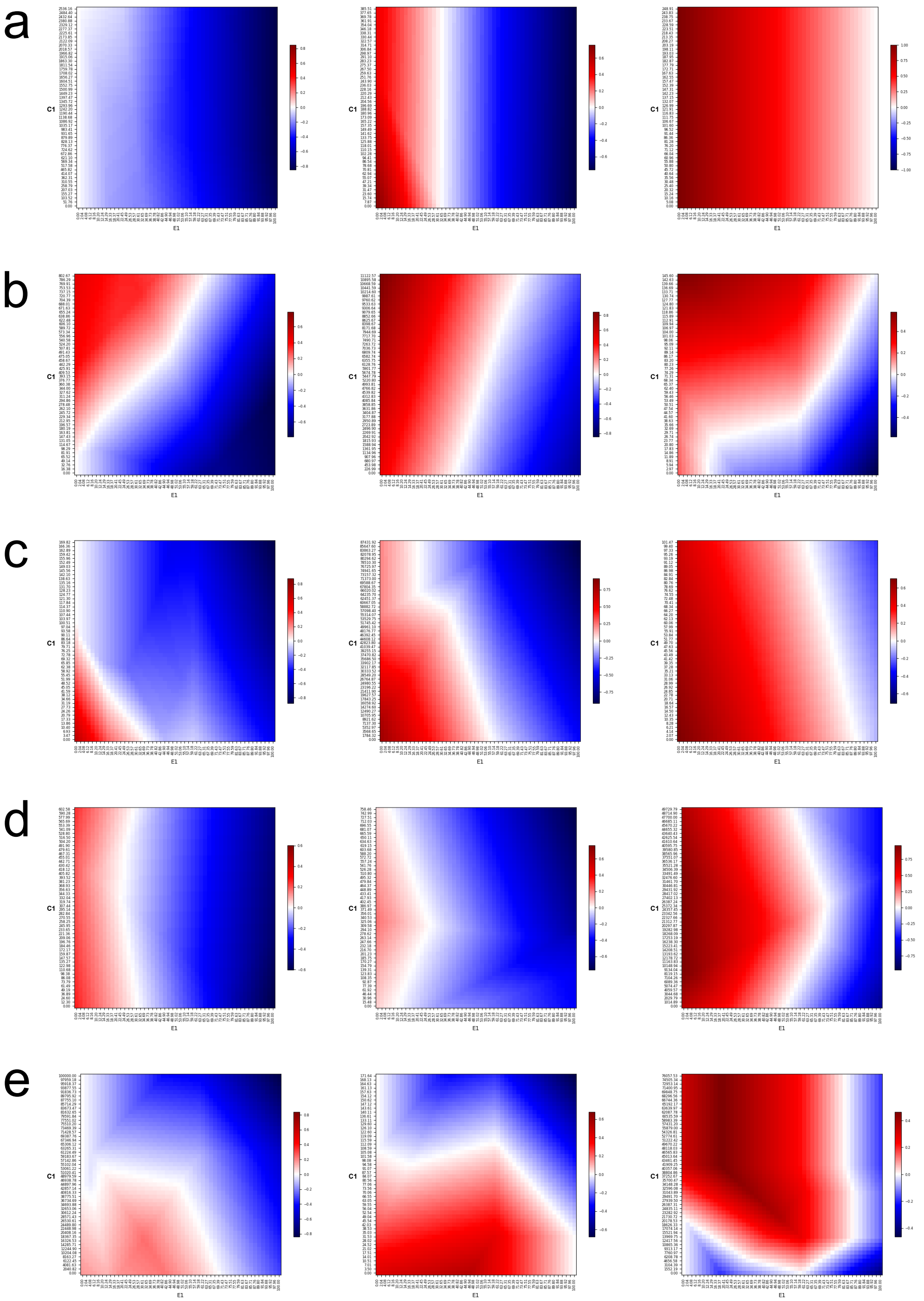


Supplementary Figure 3: Dynamic behaviour of the (2-input) regulatory network in Supplementary Figure 2 for different weight sets. Heat map x-axis and y-axis are import enzyme level $E_1$ and external nutrient concentration (given as particle number $C_1 = c_1\Omega$) input into the regulatory network, respectively. Heat map colour at $(E_1, C_1)$ shows $dE_1/d\tau$ output of the regulatory network: red = move enzyme level higher (to right), blue = move enzyme level lower (to left). It is assumed $\xi_{\sigma i}^{\mathrm{MIN}} = \xi_{\sigma i}^{\mathrm{MAX}} = 0$ in Eq. 23. Different behaviours: (a) Enzyme steady state (white region) is constant and independent of external nutrient level. (b) Enzyme steady state increases with nutrient level. (c) Enzyme steady state decreases with nutrient level. (d) Enzyme steady state has concave or convex dependence on nutrient level. (e) Enzyme steady state has hysteresis over some ranges of nutrient level.

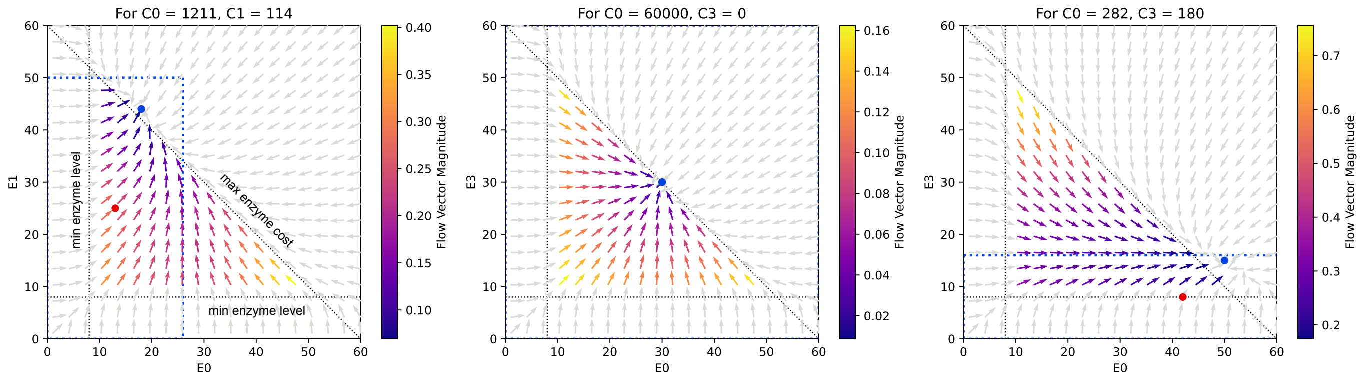


Supplementary Figure 4: Example phase plots for enzyme dynamics in a 4-input regulatory network. In each case, the regulatory network takes as input the current level of two internal import enzymes, plus the concentrations of two external nutrients that the enzymes are respectively importing. The network outputs the enzyme derivatives (shown as arrows on the plot). For each plot, the two external nutrient concentrations are held constant. Red dots indicate enzyme 'base levels'. Blue dotted boxes represent the ranges where enzyme level steady states can exist (the full plot, for the middle plot). Blue dots represent the steady state enzyme level found in simulation.

is the maximum level any single enzyme can have inside a protocell. Term $\min(e_i)$ denotes the minimum cost enzyme in the Chemical Universe (Supplementary Note 6). Term $f_i^{MAX}$ (unit C) is the maximum concentration that chemical species $i$ reaches in the reactor feed over the course of time.

2. After normalisation, entries in the input vector are further scaled up and clamped to a maximum of 1.0 by performing:

$$\widehat{E_{\sigma i}} = \min\left(10^{s^E_{\sigma i}}\overline{E_{\sigma i}},\ 1\right) \qquad \widehat{c_i} = \min\left(10^{s^c_{\sigma i}}\overline{c_i},\ 1\right) \tag{30}$$

where $s^E_{\sigma i}$ and $s^c_{\sigma i}$ are the input scaling weight *exponents* (weight vector $\mathbf{r}_\mathrm{S} = \left[s^E_{\sigma 1}, s^E_{\sigma 2}, \ldots, s^c_{\sigma 1}, s^c_{\sigma 2}, \ldots\right]$). This scaling and clamping operation allows the regulatory network to be sensitive to different order value ranges. For the model parameterisation used in this study, enzyme levels are not scaled up ($s^E_{\sigma i} = 0$). However, nutrient concentrations with their large dynamic range can have a scaling weight exponent anywhere in the continuous interval $s^c_{\sigma i} \in [0, 3]$.

3. The MLP neural network is propagated using the normalised, scaled and clamped input vector. A value in the $[0, 1]$ interval is output for the production rate of each import enzyme.

Weights in matrices $\mathbf{r}_\mathrm{B}, \mathbf{r}_\mathrm{IH}, \mathbf{r}_\mathrm{HO}$ have values in the continuous range $[-4, 4]$. The neural network has a single hidden layer of 5 nodes to keep the total number of weights low (reducing the number of weight mutations needed to change the regulatory network function).

#### Weight Values in Simulation Initial Condition

In all protocell types in the initial condition:

- Weight matrix $\mathbf{r}_\mathrm{S}$ have weights set randomly within their allowed ranges.
- Weight matrices $\mathbf{r}_\mathrm{B}, \mathbf{r}_\mathrm{IH}, \mathbf{r}_\mathrm{HO}$ have all 0 entries, making $\mathrm{NN}^i_\sigma\left([\mathbf{E}_\sigma, \mathbf{c}_\sigma]\right) = 0.5$ irrespective of the values of $\mathbf{E}_\sigma$ and $\mathbf{c}_\sigma$.

- Weights $k_{\sigma i}$ (forming weight vector $\mathbf{r}_{\mathrm{K}} = [k_{\sigma 1}, k_{\sigma 2}, \ldots]$) are set such that the 0.5 neural network output gives a desired 'base' (see below) steady state $E^*_{\sigma i}$ for each import enzyme:

$$\frac{dE_{\sigma i}}{d\tau} = k_\tau \left(0.5 - k_{\sigma i} E^*_{\sigma i}\right) = 0$$

$$0.5 - k_{\sigma i} E^*_{\sigma i} = 0$$

$$k_{\sigma i} = \frac{0.5}{E^*_{\sigma i}} \qquad \text{for } E^*_{\sigma i} \geq E_{\mathrm{MIN}} > 0 \tag{31}$$

#### Import Enzymes: Actual Levels and Base Levels

Import enzymes in a protocell type $\sigma$ have an actual level $E_{\sigma i}$ and a 'base' level $E^*_{\sigma i}$. Evolution of a protocell type changes the 'base' level of an import enzyme only (see Supplementary Note 5). This base value is used to calculate weight $k_{\sigma i}$ (via Eq. 31) of the regulatory network in Eq. 23.

At the start of simulations, regulatory networks are inactive (i.e. the neural network production term for each enzyme is 0.5 constantly). They maintain actual enzyme levels at their base values constantly: $E_{\sigma i} = E^*_{\sigma i}$. However, when a protocell type has had sufficient weight mutations made, it's regulatory network becomes responsive to inputs. From this point on, $E^*_{\sigma i}$ and $E_{\sigma i}$ can diverge as the regulatory network changes enzyme levels over short time scales in response to conditions. Importantly, $E^*_{\sigma i}$ sets the dynamic range over which $E_{\sigma i}$ can vary (see below).

If the regulatory network were to return to an inactive state, then $E_{\sigma i} = E^*_{\sigma i}$ would hold again at all times.

#### Range of Enzyme Level Dynamics and Steady States

When the base level of an import enzyme is set to $E^*_{\sigma i}$, the maximum steady state of the actual enzyme level is given when neural network output is 1.0:

$$\frac{dE_{\sigma i}}{d\tau} = k_\tau \left(1 - \frac{0.5}{E^*_{\sigma i}} E^{\max}_{\sigma i}\right) = 0$$

as $k_\tau > 0$:

$$1 - \frac{E^{\max}_{\sigma i}}{2E^*_{\sigma i}} = 0$$

$$E^{\max}_{\sigma i} = 2E^*_{\sigma i} \tag{32}$$

Conversely, the minimum steady state of the actual enzyme level is given when neural network output is 0:

$$\frac{dE_{\sigma i}}{d\tau} = k_\tau \left(0 - \frac{0.5}{E^*_{\sigma i}} E^{\min}_{\sigma i}\right) = 0$$

as $k_\tau > 0$:

$$\frac{E^{\min}_{\sigma i}}{2E^*_{\sigma i}} = 0$$

$$E^{\min}_{\sigma i} = 0 \tag{33}$$

Therefore, when unrestricted, a regulatory network can dynamically change each enzyme level in the range $0 \leq E_{\sigma i} \leq 2E^{*}_{\sigma i}$ (see blue dotted boxes in Supplementary Figure 4). In practice, the $\xi^{\mathrm{MIN}}_{\sigma i}$ and $\xi^{\mathrm{MAX}}_{\sigma i}$ terms in Eq. 23 restrict this range. The viable dynamics region for a regulatory network that controls two enzymes is illustrated as dotted lines making a triangle region in Supplementary Figure 4.

#### Speed of Regulatory Network Response

The speed at which a regulatory network can act in the valid enzyme level region can also be checked.

The maximum value of the enzyme derivative is when enzyme production is maximal (neural network output 1.0) and enzyme degradation is minimal, i.e. $E_{\sigma i} = 0$:

$$\max\left(\frac{dE_{\sigma i}}{d\tau}\right) = k_\tau \left(1 - k_{\sigma i}(0)\right) = k_\tau \tag{34}$$

Conversely, the minimum value of the enzyme derivative is when enzyme production is zero and enzyme degradation is maximal:

$$\min\left(\frac{dE_{\sigma i}}{d\tau}\right) = k_\tau \left(0 - k_{\sigma i}E^{\max}_{\sigma i}\right) = -k_{\sigma i}E^{\max}_{\sigma i}k_\tau$$

Since $E^{\max}_{\sigma i} = \frac{1}{k_{\sigma i}}$ from Eqs. (32) and (31) we have:

$$\min\left(\frac{dE_{\sigma i}}{d\tau}\right) = -k_\tau \tag{35}$$

Thus, an enzyme level can change at most by $\pm k_\tau$ units per unit time $\tau$, when in the valid enzyme level region.

When enzyme levels move outside the valid enzyme level region, terms $\xi^{\mathrm{MIN}}_{\sigma i}$ and $\xi^{\mathrm{MAX}}_{\sigma i}$ terms in Eq. 23 will cause a faster response, restoring the trajectory back into the valid region.

#### Other Remarks

- The regulatory network changes only import enzyme levels and therefore it always affects the excretion of metabolic by-products in a ratio dictated by weights $M^{\sigma}_{nb}$ of the metabolic network. In this study, the regulatory network does not change the metabolic weights $M^{\sigma}_{nb}$ of a protocell type directly.

- In this study, the regulatory network can only respond to environmental concentrations of nutrients that a protocell is importing. It is not responsive to the concentrations of other environmental chemicals.

### Supplementary Note 5 Evolution Operator

Each protocell type $\sigma$ has a phenotype consisting of:

| Metabolic weight matrix $M^{\sigma}_{nb}$ | Base levels $E^{*}_{\sigma i}$ of nutrient import enzymes | Regulatory weight matrices $\mathbf{r}_{\mathrm{S}}, \mathbf{r}_{\mathrm{B}}, \mathbf{r}_{\mathrm{IH}}, \mathbf{r}_{\mathrm{HO}}$ |
|---|---|---|
| (Floating point values) | (Integer values) | (Floating point values) |

- Each growing and dividing protocell type population produces a single copy of a variant daughter type every $\mathcal{E}_{\mathrm{div}}$ divisions, on average. This corresponds to a division event happening where one of the daughter protocells is a variant.
- The variant daughter phenotype is created by copying the parent phenotype value-for-value but introducing an error ('mutation') on each copy operation with probability $\mathcal{E}_{\mathrm{mut}}$.
- A single copy of the variant daughter protocell type is added to the reactor vessel **only if** at least 1 mutation has been made, otherwise a normal protocell division into two identical daughters is considered to have occurred.

In this work, variant daughter types always import the same nutrients and leak the same by-products as the parent type. Only minor phenotype changes are considered. Major innovations i.e speciation events involving a change of nutrient diet or leaked by-products are implemented in the platform, but are disabled for the current study.

Note that, as compared to a protocell type with a simpler machinery, a more complex protocell type could be expected to both (i) produce a variant over less division cycles and (ii) have more mutations per variant. In our model, we implement the latter effect directly. The former effect also is present to some extent indirectly: even though new variants are produced at the same frequency (every $\mathcal{E}_{\mathrm{div}}$ divisions) for all protocell types, a smaller phenotype size means that a variant is more likely to be identical to it's parents and be rejected, increasing the number of division cycles needed to create a *bona fide* variant.

#### Mutation Procedure

A value $\mathcal{X}$ in the phenotype is perturbed by drawing a small value $\epsilon$ from an exponential distribution with mean $\mu$:

$$\epsilon \sim \mathrm{Exponential}\left(\frac{1}{\mu}\right) \tag{36}$$

where mean perturbation size $\mu$ is calculated as a fraction of the maximum possible range of the value $\mathcal{X}$:

$$\mu = (\mathcal{X}_{\mathrm{MAX}} - \mathcal{X}_{\mathrm{MIN}})\mathcal{E}_{\mathrm{frac}} \tag{37}$$

The long tail of an exponential distribution permits large changes to be made infrequently.

Perturbation $\epsilon$ is added or subtracted from $\mathcal{X}$ with equal probability:

$$\mathcal{X}_{\mathrm{new}} = \mathcal{X}_{\mathrm{old}} + \epsilon \qquad \text{or} \qquad \mathcal{X}_{\mathrm{new}} = \mathcal{X}_{\mathrm{old}} - \epsilon \tag{38}$$

If the new value $\mathcal{X}_{\mathrm{new}}$ exceeds bounds $\mathcal{X}_{\mathrm{MIN}}, \mathcal{X}_{\mathrm{MAX}}$ it is clamped to the bound exceeded:

$$\mathcal{X}_{\text{new}} = \begin{cases} \mathcal{X}_{\text{MIN}} & \mathcal{X}_{\text{new}} < \mathcal{X}_{\text{MIN}} \\ \mathcal{X}_{\text{MAX}} & \mathcal{X}_{\text{new}} > \mathcal{X}_{\text{MAX}} \\ \mathcal{X}_{\text{new}} & \text{otherwise} \end{cases} \tag{39}$$

#### Mutations to Metabolic Network Weight Values

Each metabolic weight value in the phenotype selected for mutation uses:

$$\mathcal{X}_{\text{MIN}} = 0 \qquad \mathcal{X}_{\text{MAX}} = \frac{\max(v_i)}{\min(v_i)}$$

in the mutation procedure above, where $\min(v_i)$ and $\max(v_i)$ are the minimum and maximum growth values for chemicals, respectively, in the Chemical Universe (Supplementary Note 6).

**Additionally:** the new metabolic weight value must keep thermodynamic (Eq. 8) and leak (Eq. 9) constraints satisfied. If these constraints are not met, the weight value is reverted to its original value and a different perturbation is tried until a change is accepted.

#### Mutations to Enzyme Base Levels

Each enzyme base level $E^*_{\sigma i}$ in the phenotype selected for mutation uses:

$$\mathcal{X}_{\text{MIN}} = E_{\text{MIN}} \qquad \mathcal{X}_{\text{MAX}} = E_{\text{MAX}}$$

in the mutation procedure above, where $E_{\text{MAX}}$ is defined in Eq. 29. Note that the new base level $\mathcal{X}_{\text{new}}$ in Eq. 38 is rounded to the nearest integer value. A different perturbation is tried if the rounding causes $\mathcal{X}_{\text{new}} = \mathcal{X}_{\text{old}}$.

**Additionally:** the new (integer) enzyme base level must not cause the total maximum enzyme cost to be exceeded (Eq. 12); if the latter exceeded, the enzyme base level change is reverted to its original value and a different perturbation is tried until a change is accepted.

The new enzyme base level is used to set the enzyme decay weight $k_{\sigma i}$ in the regulatory network (Eq. 31), which is an element of regulatory weight matrix $\mathbf{r}_{\text{K}}$.

#### Mutations to Regulatory Network Weight Values

Each standard regulatory weight in $\mathbf{r}_{\text{B}}, \mathbf{r}_{\text{IH}}, \mathbf{r}_{\text{HO}}$ in the phenotype, which is selected for mutation uses:

$$\mathcal{X}_{\text{MIN}} = -4 \qquad \mathcal{X}_{\text{MAX}} = 4$$

while each regulatory weight in $\mathbf{r}_{\text{S}}$ uses:

$$\mathcal{X}_{\text{MIN}} = 0 \qquad \mathcal{X}_{\text{MAX}} = 3$$

in the mutation procedure above.

### Supplementary Note 6 Parameter Values for Simulations in Paper

Parameters were set with the following considerations:

- **Information about resource supply abundance is reflected in reactor vessel concentrations.** Reactor flow speed $\mu$ and total enzyme cost limit $R^{\text{enzymes}}_{\text{MAX}}$ were set so that changes in reactor feed chemical concentrations is reflected by smaller (but detectable) steady state changes in reactor vessel chemical concentrations. If $R^{\text{enzymes}}_{\text{MAX}}$ is set too high, there is nutrient scarcity in the reactor vessel because of high protocell import rates: under such conditions regulatory networks have no information about nutrient abundance.

- **Chemical number and protocell number scales are separated by orders of magnitude.** The $g$ parameter is set such that a protocell must absorb at least $\approx 1000$ nutrient particles before dividing. This ensures that protocells act more like metabolising *systems*, rather than as individual molecules undergoing reactions.

- **Computational tractability.** The number of divisions per variant daughter $\mathcal{E}_{\text{div}}$ was set relatively low such that enough variants are generated in simulation to explore the metabolic/regulatory weight space of protocell types. The reactor volume $\Omega$ was set relatively low to allow reactor vessel concentration changes to be triggered by fewer particle movement events.

| Parameter | Unit | Description | Value |
|---|---|---|---|
| **Reactor** | | | |
| $f_{\text{MAX}}$ | $C$ | Maximum concentration of nutrient in reactor feed | 200 |
| $f_{\text{MIN}}$ | $C$ | Minimum concentration of nutrient in reactor feed | 40 |
| $\mu$ | $\tau^{-1}$ | Reactor solvent flow speed | 0.02 |
| $\Omega$ | $V$ | Reactor internal volume | 500 |
| **Metabolic Network** | | | |
| $\beta$ | dimensionless | Min fraction of each nutrient growth value leaked as by-products | 0.2 |
| $R^{\text{maintain}}$ | $\upsilon\tau^{-1}$ | Self-maintenance cost | 50 |
| $R^{\text{enzymes}}_{\text{MAX}}$ | $\upsilon\tau^{-1}$ | Maximum total cost permitted for import enzymes | 60 |
| $E_{\text{MIN}}$ | $\tau^{-1}$ | Minimum permitted level of an individual import enzyme | 5 |
| $g$ | $\upsilon^{-1}$ | Protocell growth value rate to division rate converter | 0.00005 |
| **Regulatory Network** | | | |
| $k_\tau$ | dimensionless | Regulatory network timescale multiplier | 1 |
| $b_1$ | dimensionless | Push-back speed when enzyme cost exceeds $R^{\text{enzymes}}_{\text{MAX}}$ | 0.1 |
| $b_2$ | dimensionless | Boost speed when enzyme level goes below $E_{\text{MIN}}$ | 2 |
| **Evolution** | | | |
| $\mathcal{E}_{\text{div}}$ | dimensionless | Average number of divisions before variant daughter produced | 100 |
| $\mathcal{E}_{\text{mut}}$ | dimensionless | Probability to mutate phenotype parameter | 0.15 |
| $\mathcal{E}_{\text{frac}}$ | dimensionless | Fraction of max range to perturb phenotype parameter | 0.05 |

Supplementary Table 1: Model parameters.

| Case Study 1: Single Protocell Species | | |
|---|---|---|
| Chemical $n$ | Growth value $v_n$ | Growth value cost $e_n$ to maintain 1 unit of importing enzyme |
| | (unit $v$) | (unit $v$) |
| 0 | 30 | 1.0 |
| 1 | 30 | 1.0 |
| 2 | 10 | 1.0 |

| Case Study 2: Minimal Mutualism | | |
|---|---|---|
| Chemical $n$ | Growth value $v_n$ | Growth value cost $e_n$ to maintain 1 unit of importing enzyme |
| | (unit $v$) | (unit $v$) |
| 0 | 30 | 1.0 |
| 1 | 30 | 1.0 |
| 2 | 20 | 1.0 |
| 3 | 18 | 1.0 |

| Case Study 3: Binary Asymmetric Ecology | | |
|---|---|---|
| Chemical $n$ | Growth value $v_n$ | Growth value cost $e_n$ to maintain 1 unit of importing enzyme |
| | (unit $v$) | (unit $v$) |
| 0 | 40 | 1.0 |
| 1 | 40 | 1.0 |
| 2 | 20 | 1.0 |
| 3 | 2 | 1.0 |

Supplementary Table 2: Chemical Universes used in main paper Case Studies.

| | |
|---|---|
| Initial Protocells per Population | 100 |
| Initial Enzyme Levels (unit $\tau^{-1}$) | 30 |
| Initial Regulatory Network Weights | See Supplementary Note 4 |
| Initial Metabolic Network Weights | Chosen from set of initial weights that allow ecology to sustain under all forcing types |

Supplementary Table 3: Initial Conditions used in main paper Case Studies.

### Supplementary Note 7 Lineage Sizes and Variant Numbers for Simulations in Paper

| Fig. | Forcing Speed | Mode | Sim $\tau$ | p0 Lineage Size | p1 Lineage Size | Total Variants | $\tau$ per Lineage Variant |
|---|---|---|---|---|---|---|---|
| 2c | No forcing | -REG | 5M | 100 | – | 88335 | 50000 |
| | | +REG | 2M | 280 | – | 74089 | 7143 |
| 2d | Slow | -REG | 5M | 204 | – | 52083 | 24510 |
| | | +REG | 5M | 782 | – | 108534 | 6394 |
| 2e | Medium | -REG | 5M | 209 | – | 52211 | 23923 |
| | | +REG | 5M | 940 | – | 105500 | 5319 |
| 2f | Fast | -REG | 5M | 199 | – | 52585 | 25126 |
| | | +REG | 5M | 1005 | – | 108231 | 4975 |
| 3b | No forcing | -REG | 5M | 158 | 176 | 111585 | 29940 |
| | | +REG | 2M | 287 | 350 | 87958 | 6279 |
| 3c | Slow | -REG | 5M | 240 | 306 | 64751 | 18315 |
| | | +REG | 5M | 850 | 841 | 137522 | 5914 |
| 3d | Medium | -REG | 5M | 277 | 268 | 63783 | 18349 |
| | | +REG | 5M | 861 | 886 | 130702 | 5724 |
| 3e | Fast | -REG | 5M | 263 | 281 | 64316 | 18382 |
| | | +REG | 5M | 840 | 960 | 130649 | 5556 |

Supplementary Table 4: Evolutionary lineage sizes for simulations in Figure 2 and Figure 3 of main paper. The 'Sim $\tau$' column shows total simulation time (M = million time steps). The last column shows the mean number of time steps that elapses between birth of significant variants that make up the 'trunk' of the evolutionary lineage tree(s). This places a limit on the speed of phylogenetic adaptation.

#### Supplementary Note 8 Computational Optimisations

The model includes protocell metabolism (with regulatory control), ecology and evolution. Thus, simulations typically require considerable computational resources. In the code design, significant steps were taken to reduce computational costs:

- Gillespie SSA optimisations:
  - A relatively small reactor volume ($\Omega = 500$ particles per concentration unit) was used such that concentration levels could be changed with fewer transitions firing.
  - The core SSA function ran as machine code, JIT compiled by the Python NUMBA library
  - The propensity vector was selectively updated: only rates affected by the last firing event were re-calculated
  - Total system propensity value was maintained as a diff (not completely re-calculated after each firing transition)
- Neural network optimisations
  - Caching of neural network calculations was used to allow a regulatory network to be forward-propagated only once each time the input vector changed.
  - Linear sigmoid transfer functions were used to avoid more expensive $e^x$ calculations.

With these optimisations, a $5 \times 10^6$ time unit simulation with regulatory networks enabled typically:

- Fires from $3.5$ to $5.5 \times 10^{10}$ Gillespie events.
- Takes 24 hours wall time on a modern computer. This equates to $3500$ time units simulated per minute wall time.
- Generates approx 300k files = 1Gb of compressed data = 0.06 files per time unit.

[S6] Ricky T. Q. Chen, Yulia Rubanova, Jesse Bettencourt, and David Duvenaud. Neural Ordinary Differential Equations. *arXiv Preprint*, 1806.07366, 2018.
